# RNA helicase DDX6 in P-bodies is essential for the assembly of stress granules

**DOI:** 10.1101/2021.09.24.461736

**Authors:** Vladimir Majerciak, Tongqing Zhou, Zhi-Ming Zheng

## Abstract

Two prominent cytoplasmic RNA granules, ubiquitous RNA-processing bodies (PB) and inducible stress granules (SG), regulate storage of translationally arrested mRNAs and are intimately related. In this study, we found the dependence of SG formation on PB in the cells under arsenite (ARS) stress, but not the other way around. GW182, 4E-T and DDX6 essential for PB formation differentially affect SG formation in the cells under ARS stress, with DDX6 being the most prominent. The cells with DDX6 deficiency display irregular shape of SG which could be rescued by ectopic wt DDX6, but not its helicase mutant E247A DDX6, which induces SG in the cells without stress, indicating that DDX6 helicase activity is essential for PB, but suppressive for SG. DDX6’s dual roles are independent of DDX6 interactors EDC3, CNOT1, and PAT1B. This study provides a conceptual advance of how DDX6 involves in the biogenesis of PB and SG.

**Highlights:**

- PB act as the seeds for SG nucleation
- PB components colocalize with SG
- DDX6 plays a critical role in biogenesis of both PB and SG
- DDX6 helicase activity prevents SG formation in normal cells

## Introduction

Cytoplasmic membrane-less RNA-processing bodies (PB) and stress granules (SG) are two of the best-studied, common RNA granules. Both PB and SG are made up of translationally arrested mRNAs in complex with numerous RNA-binding proteins (RBP) and thus are associated with translational control in eukaryotic cells (Riggs et al., 2020). Despite the functional overlaps, PB and SG have profound differences in their composition, assembly, and dynamics.

The smaller PB occur in every cell under physiological conditions and are ubiquitous from yeast to human (Kedersha and Anderson, 2007). PB, free from 40S ribosomal subunits, are derived from the condensation of various ribonucleoprotein (RNP) complexes, including ∼125 RBP. Two-thirds of PB-associated RBP are functionally linked to mRNA repression, RNA decay, miRNA-mediated silencing, RNA decapping, and non-sense mediated RNA decay (NMD) (Hubstenberger et al., 2017; Youn et al., 2018). However, it has become increasingly clear that PB are not the sites of mRNA degradation (Ayache et al., 2015; Eulalio et al., 2007; Horvathova et al., 2017; Parker and Sheth, 2007; Sharma et al., 2019; Sheth and Parker, 2003) and mainly function as the sites of translational repression of AU-rich, nonoptimal codon-enriched mRNAs encoding the proteins with regulatory functions (Courel et al., 2019; Matheny et al., 2019). The mRNA translational repression in PB is independent of miRNA-mediated silencing (Eulalio et al., 2007; Sharma et al., 2019). In fact, mRNAs in the PB are actually protected from decay and become translatable after release from PB (Aizer et al., 2014; Brengues et al., 2005; Hubstenberger et al., 2017; Wilbertz et al., 2019). Interestingly, the mRNAs encoding house-keeping proteins and non-coding lncRNAs are generally excluded from PB (Hubstenberger et al., 2017). DCP1A, DCP2, GW182, DDX6, 4E-T, and EDC3 are PB-specific protein markers commonly used to visualize the PB in the cells (Ayache et al., 2015; Kedersha et al., 2008; Minshall et al., 2009; Sharma et al., 2019).

The larger SG contain 40S ribosomal subunits and appear in the cells only under stress conditions (Hofmann et al., 2021). The cells under both biological (hypoxia, nutrients deprivation or viral infection) and environmental (changes in temperature or osmotic pressure) stress or exposure to certain chemicals (sodium arsenite or ARS, chemotherapy compounds, etc.) can be induced to the formation of SG due to rapid increase of the translationally arrested mRNAs released from polysomes (Bounedjah et al., 2012; Kedersha et al., 1999; Kwon et al., 2007; Sharma et al., 2017). Different types of stress induce phosphorylation of an essential translation initiation factor eIF2α at Ser51 position (Kedersha et al., 1999; Lu et al., 1999) by activation of four different eIF2α kinases (GCN2, PKR, PERK, and HRI) (Donnelly et al., 2013; Harding et al., 2000; McEwen et al., 2005; Srivastava et al., 1998; Wek et al., 1995). Heterotrimeric eIF2, which is composed of eIF2α, eIF2β and eIF2γ, binds GTP and the initiator methionine-tRNA (tRNAi^Met^) as an eIF2-GTP-tRNAi^Met^ complex and loads the tRNAi^Met^ onto the small 40S ribosomal subunit by hydrolysis of eIF2-GTP to eIF2-GDP. Phosphorylation of eIF2α leads eIF2 stably bind to eIF2B, a guanine exchange factor and sequester the eIF2B activity, thus inhibiting the recycling of eIF2-GDP to eIF2-GTP by eIF2B and consequently blocking the formation of preinitiation complexes and protein translation (Gray and Wickens, 1998; Jackson et al., 2010; Kedersha et al., 2002). Resulting translationally stalled mRNAs with bound small ribosomal subunits are then recognized by G3BPs and TIA-1 nucleators for SG assembly and accumulated or stored in the SG (Gilks et al., 2004; Kedersha et al., 2000; Kedersha et al., 2016; Kedersha et al., 1999; Tourrière et al., 2003). The chemical inhibitor antagonizing phospho-eIF2 effects on eIF2B restores translation and causes SG disassembly (Sidrauski et al., 2015). However, in some cases a type II SG can be formed independently of eIF2α phosphorylation (Dang et al., 2006; Emara et al., 2012). Conceivably, most of the SG protein components are the proteins involved in translation initiation and regulation. To date, thousands of the different mRNA species with relative longer coding and UTR regions and ∼411 RBP have been identified in purified SG from the cells under stress (Banani et al., 2016; Matheny et al., 2019). However, the mRNAs within the SG are not degraded and become translatable after recovery from stress (Wilbertz et al., 2019). An ATR4 reporter RNA was found being translatable even within the SG itself (Mateju et al., 2020). RNA-binding proteins TIA-1, G3BP1, and PABPC1 are SG-specific markers useful for visualization of the SG in the cells under various stress conditions.

PB and SG are dynamically linked sites of RNP remodeling and share some of their components (Kedersha et al., 2005; Matheny et al., 2019). The mechanisms of how PB in cells under physiological conditions and SG in the cells under stress are formed are only partially disclosed (Ivanov et al., 2019). Previous observations indicated that PB are commonly increased in number and size in the cells during stress and often appear in close physical proximity with SG, but however, their intimate relationship is not fully understood. Both PB and SG are enriched in RBP containing intrinsically disordered regions (IRD) or prion-like domains (PrD) (Jain and Parker, 2013; Protter et al., 2018; Sanders et al., 2020). These RBP at high local concentration mediate multivalent protein-RNA, protein-protein, and RNA-RNA interactions that lead to liquid-liquid phase separation (LLPS) (Van Treeck and Parker, 2018). What drives their high local concentration required for LLPS is unknown. Several proteins were found to be essential for PB formation in normal conditions, including DDX6, 4E-T, LSM14A, GW182 (Ayache et al., 2015; Minshall et al., 2009; Sharma et al., 2019). SG formation is mechanistically better understood as described above. Their formation is driven by a high concentration of stalled preinitiation complexes recognized by SG nucleating RBP, such as G3BP1, TIA-1 and others, which are capable of binding both RNA and small ribosomal subunits and thus mediate multivalent interactions to facilitate SG condensation and recruitment of additional factors (Gilks et al., 2004; Kedersha et al., 2000; Tourrière et al., 2003). These multivalent nucleation interactions cause LLPS to form SG and can be regulated by a G3BP-binding CAPRIN1 to promote or another G3BP-binding USP10 to prevent SG formation (Kedersha et al., 2016). Microscopic studies showed that both granules represent highly dynamic structures, rapidly exchanging their components with the surrounding environment (Kedersha et al., 2005; Mollet et al., 2008). This observation indicates a constant, ATP-dependent rearrangement of RNP for SG formation (Jain and Parker, 2013; Tauber et al., 2020). Several ATP-dependent RNA helicases are enriched in both PB and SG (Hubstenberger et al., 2017; Markmiller et al., 2018; Tauber et al., 2020; Youn et al., 2018). Their ability to modulate the RNP composition was proposed to contribute to PB and SG dynamic nature (Hilliker, 2012; Tauber et al., 2020).

DEAD-box helicase 6 (DDX6/p54/Rck and its orthologues Dhh1p in *S. cerevisiae*, Me31B in *D. melanogaster*, CGH-1 in *C. elegans*, and Xp54 in *X. laevis*) is one of the ATP-dependent, cellular RNA helicases containing a conserved Asp-Glu-Ala-Asp motif (DEAD-box) and other high levels of sequence and structure conservations (Buchan et al., 2008; Cordin et al., 2006; Linder and Jankowsky, 2011; Ostareck et al., 2014; Weston and Sommerville, 2006). DDX6 consists of an N-terminal and a C-terminal domain for interactions with various factors for specific functions and two RecA-like domains harboring both ATP-binding/ATPase and RNA-binding/helicase activities (Sarkar and Ghosh, 2016). DDX6, abundantly expressed in cells, is highly interactive with RNAs and almost half of all PB proteins (Ayache et al., 2015; Ernoult-Lange et al., 2012; Hubstenberger et al., 2017). Thus, DDX6 has been found to be essential for PB formation and retaining PB identity during stress (Serman et al., 2007). Interestingly, while ATPase activity is essential for PB assembly, this activity of DDX6 does not have any effect on the existing PB (Minshall et al., 2009). The top interactors of DDX6 include members of mRNA decay and translational repression complexes (Ayache et al., 2015). The C-terminal RecA-like domain of DDX6 mediates the most of DDX6 protein-protein interactions, such as interactions with PAT1B, CNOT1 and 4E-T in the regulation of PB formation (Ayache et al., 2015; Ozgur et al., 2015; Tritschler et al., 2009). Although DDX6 is a cofactor of XRN1 5’−3’ exonuclease and its interaction with PAT1B affects RNA 3’ to 5’ degradation, recent reports indicate that there is no RNA decay or degradation within the PB (Aizer et al., 2014; Brengues et al., 2005; Hubstenberger et al., 2017; Wilbertz et al., 2019). While these protein-protein and protein-RNA interactions seem sufficient for DDX6 to act as a central node, it becomes increasingly clear that DDX6 RNA-helicase activity appears essential for de novo PB formation (Di Stefano et al., 2019; Minshall et al., 2009).

Up to date, the exact intimate nature between PB and SGs is yet to be understood, although both contain translationally repressed RBP and shared exchangeable components (Kedersha et al., 2005). In this study, we investigated the contribution of PB to SG formation in HeLa cells in response to ARS treatment. We found that most SG sprout from existing PB and are enriched in the expected PB components. RNAi screen led us to identify several PB components which are required for SG formation, but DDX6 was found being the major factor essential for PB formation in cells without or with the ARS treatment. Interestingly, we found that loss of DDX6 was detrimental not only for PB but also for SG biogenesis. We found that the dual activities of DDX6 in the biogenesis of both PB and SG were independent of its binding partners and the cells deficient of DDX6 could be restored to form typical SG by the ectopic expression of the exogenous DDX6. On the contrary, expression of the helicase-deficient DDX6 mutant led to the formation of SG-like granules even in the absence of stress. In summary, we identified an essential role of PB in SG biogenesis, with DDX6 playing a central role.

## Results

### Kinetics of stress granule formation in cells

Previous observations suggested a close topological, biochemical, and functional connection between the PB and SG (Kedersha et al., 2005; Riggs et al., 2020; Youn et al., 2018). However, the role of PB in SG formation remains elusive. To determine whether any contribution of PB components to SG formation, we first performed a time-course study on SG biogenesis in HeLa cells plated on the cover glass after exposing to ARS (0.5 mM), a well-studied SG inducer, for 0, 5, 10, 15, 20, and 30 min and following by staining for GW182, a renown marker of PB, and TIA-1, a renown marker for SG. As shown in **Figure 1**, each HeLa cell at both time 0 and 5 min of ARS induction exhibited ∼4-5 large and ∼10-15 smaller rounded GW182-positive PB as previously reported (Sharma et al., 2019), but no detectable SG. At this stage, TIA-1 was stained as a diffused cytoplasmic protein with some nuclear appearance. By 10 min, but more profoundly by 15 min, we started to observe the appearance of granular TIA-1 staining in the cytoplasm of ARS-treated cells, accompanied by an increased number of small PB. Interestingly, most newly formed SG appeared overlapping with or in close physical proximity to PB, with mature SG being often surrounded by multiple PB (see zoom in 30 min time point), although most but not all PB were associated with the SG. Together, these data strongly suggest that SG are spatially intricately linked to preexisting PB, which may act as the seeds to initiate SG formation.

**Figure 1.**
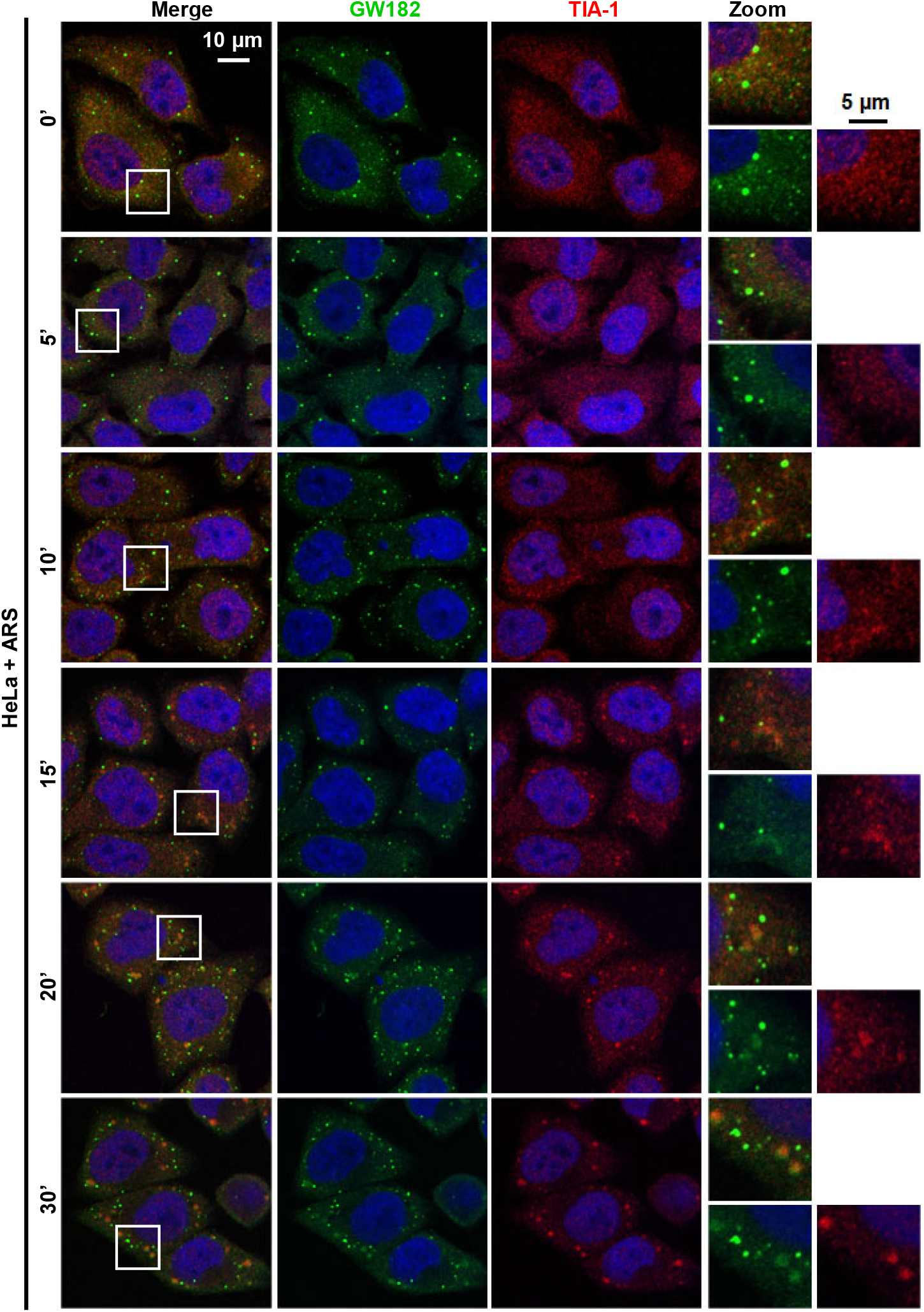
PB act as the seeds for stress granules (SG) formation during ARS stress. HeLa cells were grown on the cover glass and 24 h later exposed to 0.5 mM ARS. The cells at time 0, 10, 15, 20, and 30 min (labeled on the left) were fixed and stained for PB with anti-GW182 and SG with anti-TIA-1 antibodies.

### Identification of PB components in the newly formed SG

A close examination of GW182 staining in ARS-treated cells showed that ARS not only induces SG but also affects PB structure. As shown in **Figure 1**, the PB became larger and more distinct upon the stress (compare GW182 staining at 0 and 30 min). In addition, we observed GW182 accumulation in newly formed SG as measured by the distribution of GW182 signal intensity across the PB and connected SG. As shown in **Figure 2A**, GW182 signal intensity within the PB was ∼4-5 fold above or ∼2-3 fold within the G3BP1^+^ SG above the background staining in the surrounding cytoplasm. Because the GW182 signal within the SG was tightly overlapped with G3PB1 staining, this data indicates the accumulation of GW182 in the newly formed SG. We next asked if other PB components also accumulate in SG. We examined two additional, commonly used PB-specific markers, DDX6 and DCP1A (**Figure 2B-2C**). We found that DDX6, a DEAD-box RNA helicase, also colocalized with G3BP1 in the SG beside its intense staining within the PB (**Figure 2B**). In contrast, DCP1A, a member of the RNA decapping complex, remained within the PB even during the stress conditions (**Figure 2C**). We occasionally found the overlap staining between DCP1A and G3BP1 at the boundaries between PB and SG, indicating the existence of a peripheral, close physical connection from some PB to SG. These data suggest that some PB components, such as GW182 and DDX6, are colocalized with newly formed SG, while DCP1A remains physically associated predominantly with PB. The data is consistent with the previously reported partial overlap in PB and SG proteomes (Youn et al., 2018).

**Figure 2.**
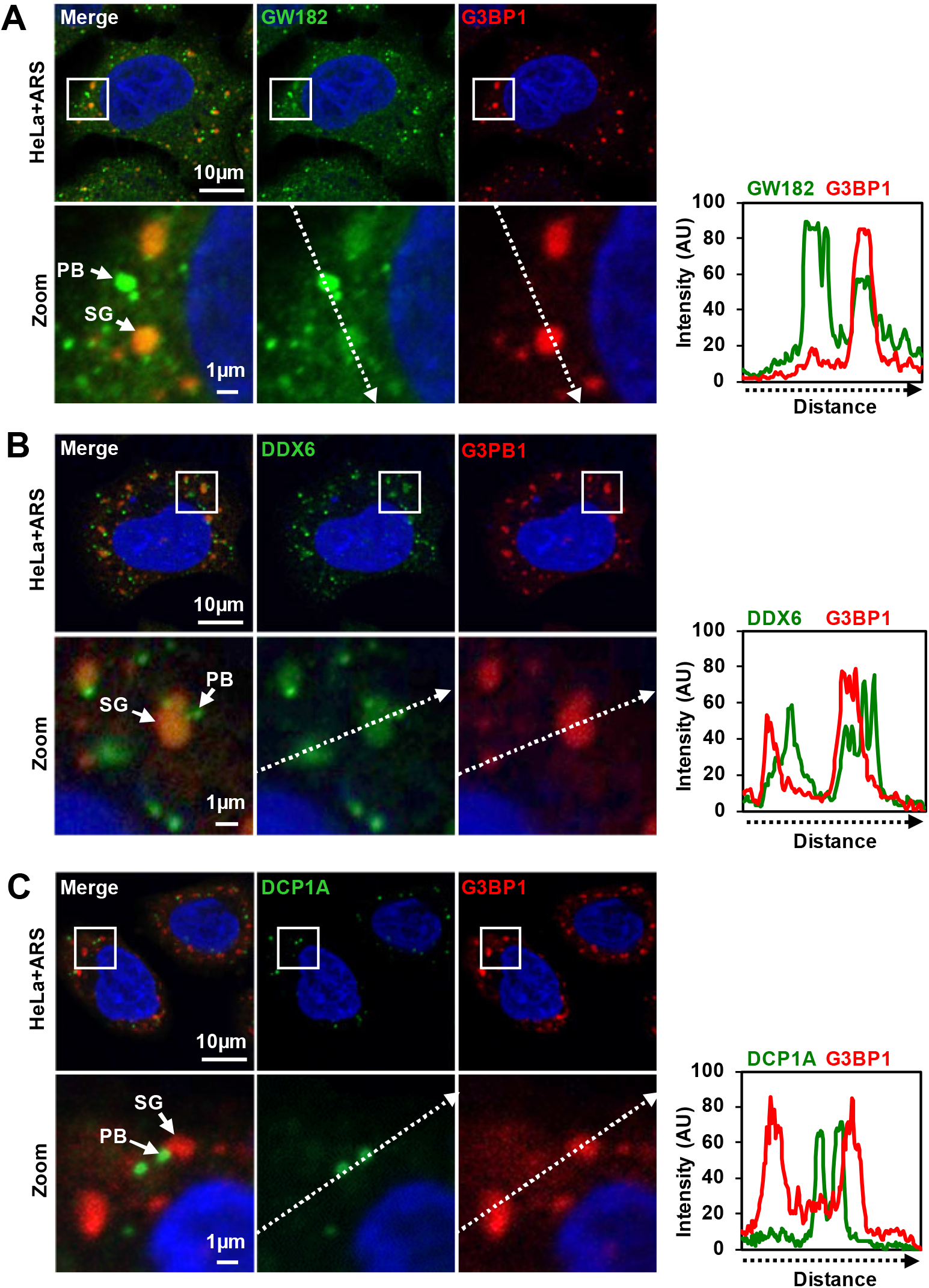
Localization of PB components in SG. HeLa cells were grown on the cover glass for 24 h and treated with 0.5 mM ARS for 30 min before staining with anti-GW182 (A), anti-DDX6 (B), and anti-DCP1A (C) antibodies for PB and anti-G3PB1 for SG. The graphs on the right show signal intensity measured by ImageJ across the image section marked by each dashed arrow.

### Roles of PB components in SG formation

Based on the previously observed topological and proteomic overlaps between PB and SG, we asked if any specific PB component plays an active role in SG biogenesis. To answer this question, we performed various siRNA knockdown (KD) experiments in HeLa cells using gene-specific siRNAs targeting major PB components, including GW182 (TNRC6A) (Sharma et al., 2019), EDC3 (Tritschler et al., 2009), and DDX6 (Ayache et al., 2015; Minshall et al., 2009). The cells treated with non-targeting siRNA (siNT) served as a negative control, and the cells treated with the siRNA targeting SG nucleating protein TIA-1 served as a positive control. Western blot confirmed the KD efficiency of GW182 by ∼76%, EDC3 by ∼81%, and DDX6 by ∼77% when compared to the cells treated with siNT (**Figure 3A**). TIA-1 was consistently reduced only ∼50% in HeLa cells despite using a pool of validated TIA-1 siRNAs in multiple repeats, suggesting a possible long half-life and high abundance of TIA-1 protein *in vivo*. After 48 h KD of individual gene expression, the cells were exposed to ARS (0.5 mM) treatment for 30 min to induce SG marked by G3PB1 staining. As shown in **Figure 3B**, the control cells treated with siNT exhibited numerous typical SG, indicating that siRNA-treatment per se had no detrimental effect on SG formation. As expected, the cells with ∼50% KD of TIA-1 displayed a reduced number of SG but had no effect on PB formation. KD of EDC3, which is not required for PB formation (Ayache et al., 2015) (**Figure S1**) did not prevent the formation of either SG or PB (**Figure 3B**). Interestingly, KD of GW182 (TNRC6A) or DDX6 which prevents the formation of PB in normal cells (**Figure S1**) (Ayache et al., 2015; Minshall et al., 2009; Sharma et al., 2019) led to either reduction of the SG number (GW182 KD) or change of the SG morphology (DDX6 KD) in ARS-treated cells. In addition, ARS-induced SG in the cells with GW182 KD or DDX6 KD could be stained sometimes by another PB marker DCP1A. We also noticed in the cells with DDX6 KD that the irregular shape of G3BP1^+^ SG overlapped with diffused, weak DCP1A signal (**Figure 3B**). Altogether, these data suggest that GW182 or DDX6, which is essential for PB formation in normal cells under physiological conditions, is dispensable for ARS-induced PB formation in the cells under stress conditions, and further indicate that PB biogenesis in the cells during the stress differs from the one under normal cell conditions as previously proposed (Ohn et al., 2008).

**Figure 3.**
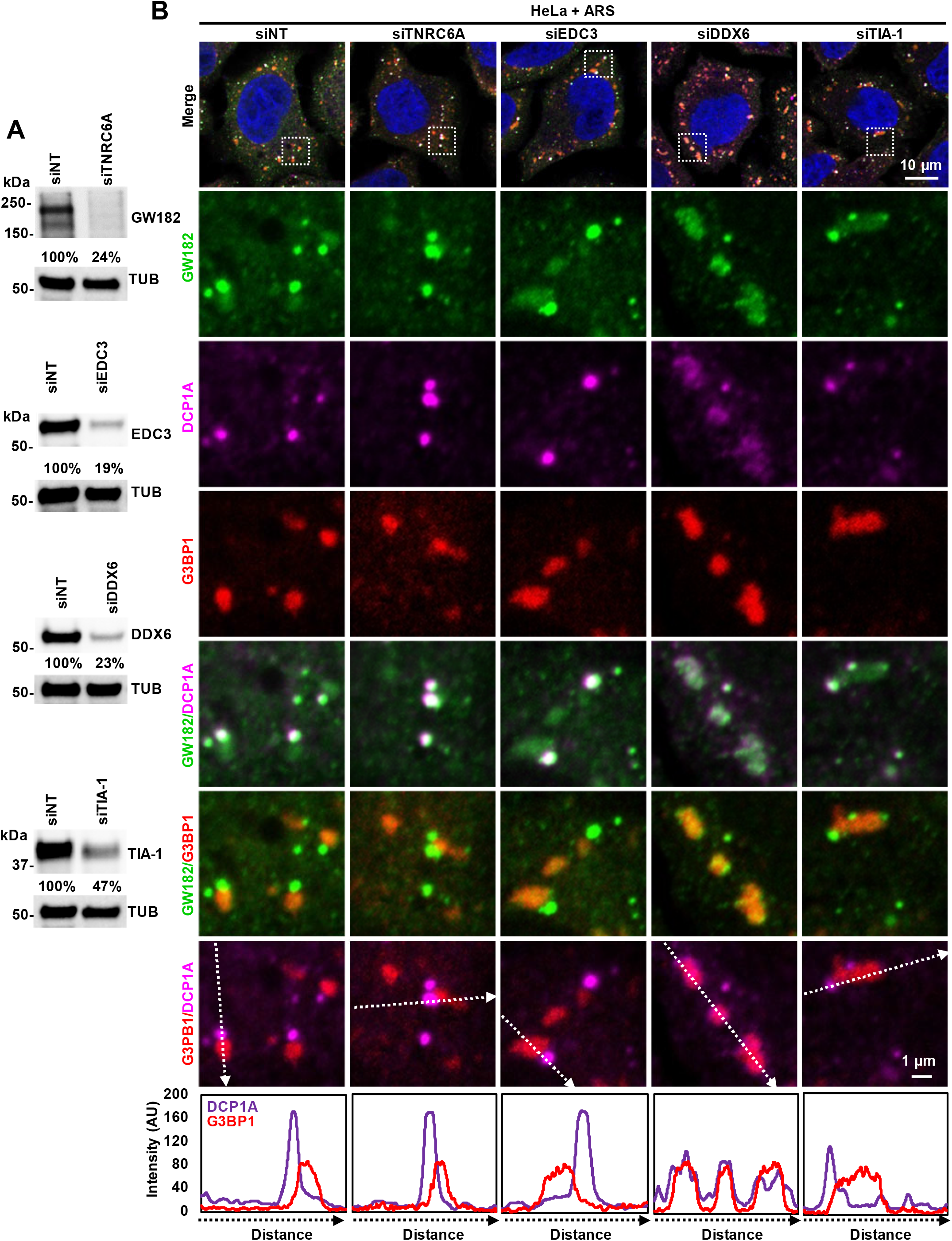
PB components play an important role in stress granules formation. HeLa cells were treated twice with control non-targeting (siNT) or gene-specific siRNAs in a 24 h interval for 48 h before ARS treatment (0.5 mM, 30 min). (A) Knockdown (KD) efficiency of a specific gene expression was determined by Western blotting analysis by comparison to the protein level of NT siRNA-treated cells after the normalization to β-tubulin (TUB). (B) The cells were then fixed and stained with an anti-GW182 human serum (green) and an anti-DCP1A for PB (magenta), and an anti-G3BP1 antibody for SG (red), respectively. Below are the signal intensity plots measured by ImageJ for DCP1A and G3BP1 across the image section marked by a dashed arrow.

To confirm this observation, we used series of *z*-stack images to generate the cell 3D models and investigated the extent of PB and SG interactions in the ARS-treated HeLa cells with normal or reduced DDX6 expression by siRNA KD. As expected in siNT-treated control cells, we observed well-defined GW182^+^ PB and G3BP1^+^ SG and retained the expected shape (**Figure 4A**). Both granules were distinguishable from each other and remained their close proximity or sometimes overlapped. In contrast, the DDX6 KD cells exhibited structurally poorly defined granules positive for both GW182 and G3PB1 staining and loss of their unique identity (**Figure 4B**). Further analysis of 50 cells in each group revealed on average that the cells treated with siNT had ∼40.5 SG per cell, but the cells treated with siDDX6 showed ∼54 irregular SG per cell (**Figure 4C**). Together these data suggest that DDX6 KD had a profound detrimental effect on the biogenesis of both PB and SG in the cells under ARS stress.

**Figure 4.**
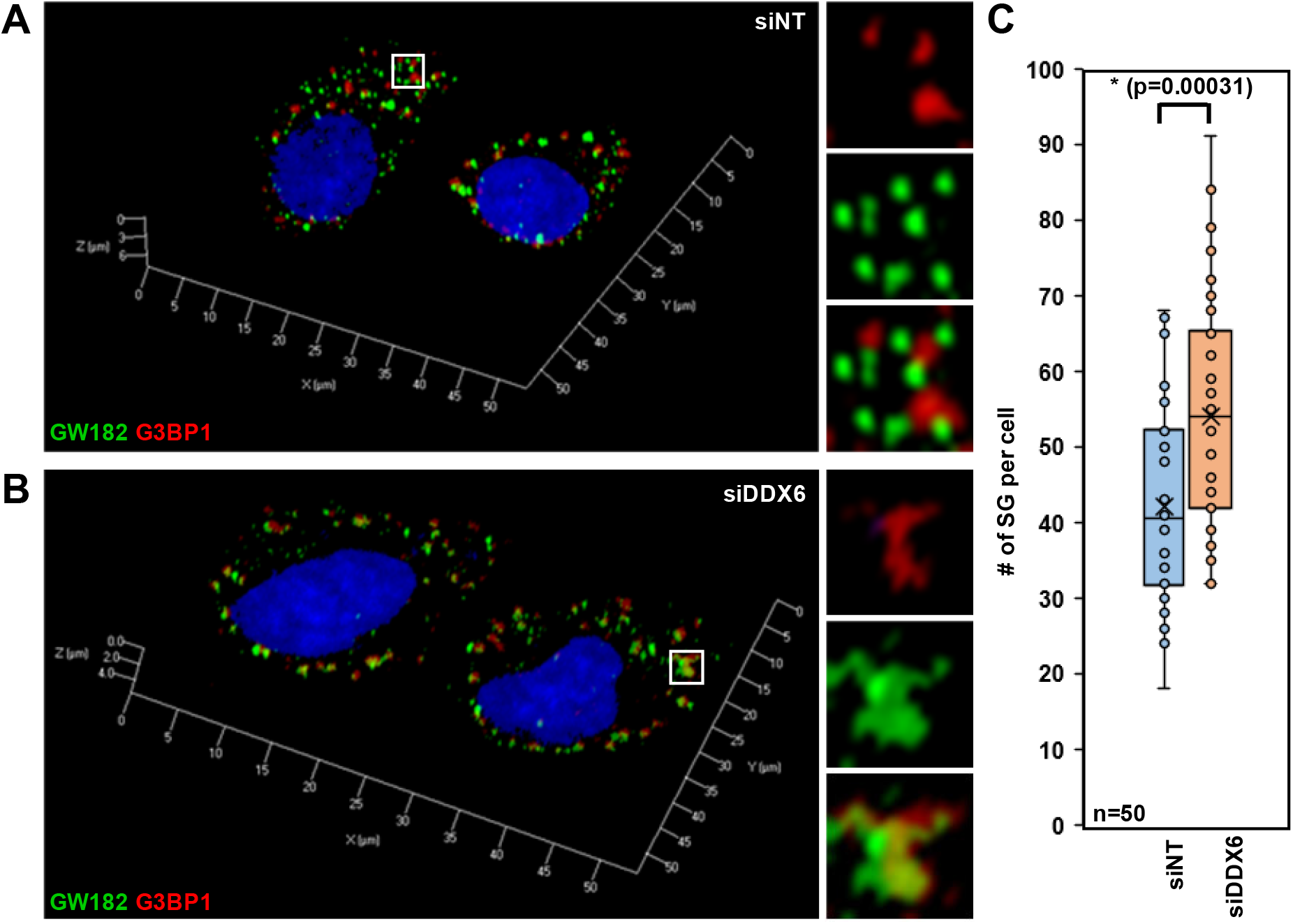
Knockdown (KD) of DDX6 expression led to aberrant SG formation. (A-B) 3D reconstruction of PB and SG in HeLa cells transfected with non-targeting (siNT, A) and DDX6 targeting (siDDX6, B) siRNAs after ARS (0.5 mM, 30 min) treatment. PB were visualized with GW182 and SG with G3BP1 staining. The *z*-stack images were collected by confocal microscopy and converted to 3D models using the maximum intensity mode. (C) G3BP1-positive SG per cell in HeLa cells treated with non-targeting (siNT) and DDX6-targeting (siDDX6) siRNAs were averaged from 50 cells in A and B. The p-value was calculated using two-tailed Student’s *t*-test.

### DDX6 activity in SG biogenesis is independent of its interaction with EDC3, PAT1B, and CNOT1, but partially on 4E-T

To examine whether DDX6-interacting partners PAT1B, CNOT1, and 4E-T are functionally required for DDX6-mediated SG biogenesis in addition to their reported role in PB formation (Kamenska et al., 2016; Luo et al., 2018; Ozgur et al., 2015), we knocked down the expression of PAT1B, CNOT1, and 4E-T in HeLa cells by gene-specific siRNAs, with the non-targeting siRNA siNT described above serving as a control (**Figure 5**). Consistent with other previous reports (Kamenska et al., 2016), KD of PAT1B and CNOT1 did not affect the formation of DCP1A^+^ PB (**Figure S2**) but did so by KD of 4E-T, which is essential for PB formation as reported (Ayache et al., 2015; Marnef et al., 2012; Ohn et al., 2008; Ozgur et al., 2015). When these cells with KD of each gene expression of the three were exposed to ARS (0.5 mM) for 30 min, we observed none of their deficiency in the cells could affect the formation of G3BP1^+^ SG (**Figure 5B**) despite 4E-T KD blocking the formation of DCP1A^+^ PB and slightly reducing number and size of SG in the cells (**Figure 5B**).

**Figure 5.**
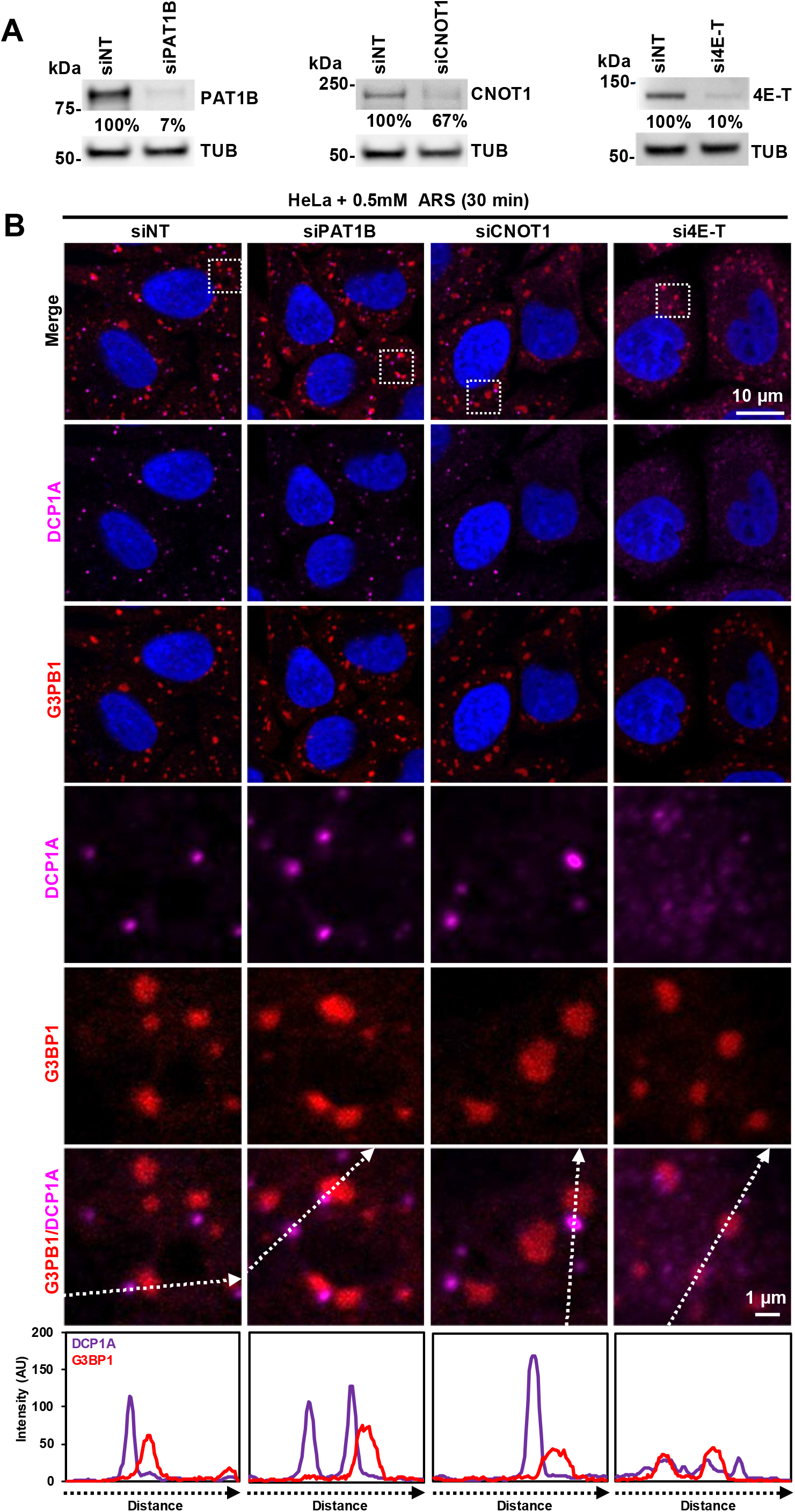
DDX6 interactors PAT1B, CNOT1 and 4E-T do not contribute to PB and SG biogenesis. HeLa cells were treated twice with siRNA targeting PAT1B, CNOT1, or 4E-T in a 24 h interval and induced with 0.5 mM ARS treatment for 30 min. (A) Knockdown efficiency by siRNA to the indicated protein expression was examined by Western blotting using a corresponding antibody, with β-tubulin (TUB) serving as an internal loading control. (B) The fixed HeLa cells were stained with an anti-DCP1A antibody for PB and an anti-G3BP1 antibody for SG. Below are the signal intensity plots measured by ImageJ for DCP1A and G3BP1 across the image section marked by a dashed arrow.

Considering that the SG in the 4E-T KD cells lack visible DCP1A^+^ PB and the DCP1A is nonessential for PB formation (Sharma et al., 2019), we wished to confirm this observation by DDX6 staining (**Figure S3**). The cells treated with control siRNA siNT exhibited typical DDX6^+^ PB and typical G3BP1^+^ SG induced by ARS (**Figure S3A**). On the contrary, we found that the cells lacking 4E-T expression did exhibit fewer distinct DDX6^+^ PB (**Figure S3B**), which were morphologically different from the diffused DCP1A staining in the 4E-T KD cells (**Figure 5B**), confirming its requirement for PB formation under normal cell conditions (**Figure S2, Figure S3B**) and a potential role of 4E-T in recruiting DCP1A to PB (Ferraiuolo et al., 2005). However, after ARS induction, the 4E-T KD cells appeared numerous distinct DDX6^+^ PB (**Figure S3B**). Notably, most of these DDX6^+^ PB were either overlapped or in close proximity with irregular SG (**Figure S3B**). These data clearly showed the PB component DDX6 in HeLa cells is required for proper SG biogenesis during the chemical stress, although 4E-T might play a minor role.

### The helicase activity of DDX6 is essential in biogenesis of SG in the cells under stress

Considering that the helicase activity of RNA helicases plays a role in the remodeling of RNP complexes and the ATPase activity of DDX6 is essential for PB assembly (Minshall et al., 2009), we next investigated whether this ATPase activity of DDX6 is also important for SG formation in the cells under ARS stress. Subsequently, we constructed a wild-type (wt) DDX6-myc-FLAG and its helicase-deficient mutant by converting glutamic acid residue 247 to alanine (E247A) in DEAD-box motif required for ATPase activity for functional studies in the formation of PB and SG in HeLa cells (**Figure 6A**). The E274A mutation eliminates critical ATP-binding/hydrolysis interaction in the DDX6 DEAD-box motif (**Figure 6B**). The presence of the epitope tags (myc-FLAG) on the ectopically expressed DDX6 allowed it to be distinguishable from the endogenous DDX6. To test if the ectopically expressed DDX6 behaves functionally similar to endogenous DDX6, we expressed the ectopic wt DDX6 and its E247A mutant protein in HeLa cells without or with ARS treatment and stained the cells for PB formation with an anti-myc (**Figure S4**) or anti-FLAG antibody (**Figure 6C-6D, Figure S5**) to detect ectopic DDX6 and an anti-DDX6 antibody or an anti-GW182 serum to detect both endogenous and ectopic DDX6 proteins (**Figure 6C-6D, Figure S5**), and for SG formation with anti-TIA-1, anti-PABPC1 and anti-G3BP1 antibodies (**Figure S4, Figure 7**).

**Figure 6.**
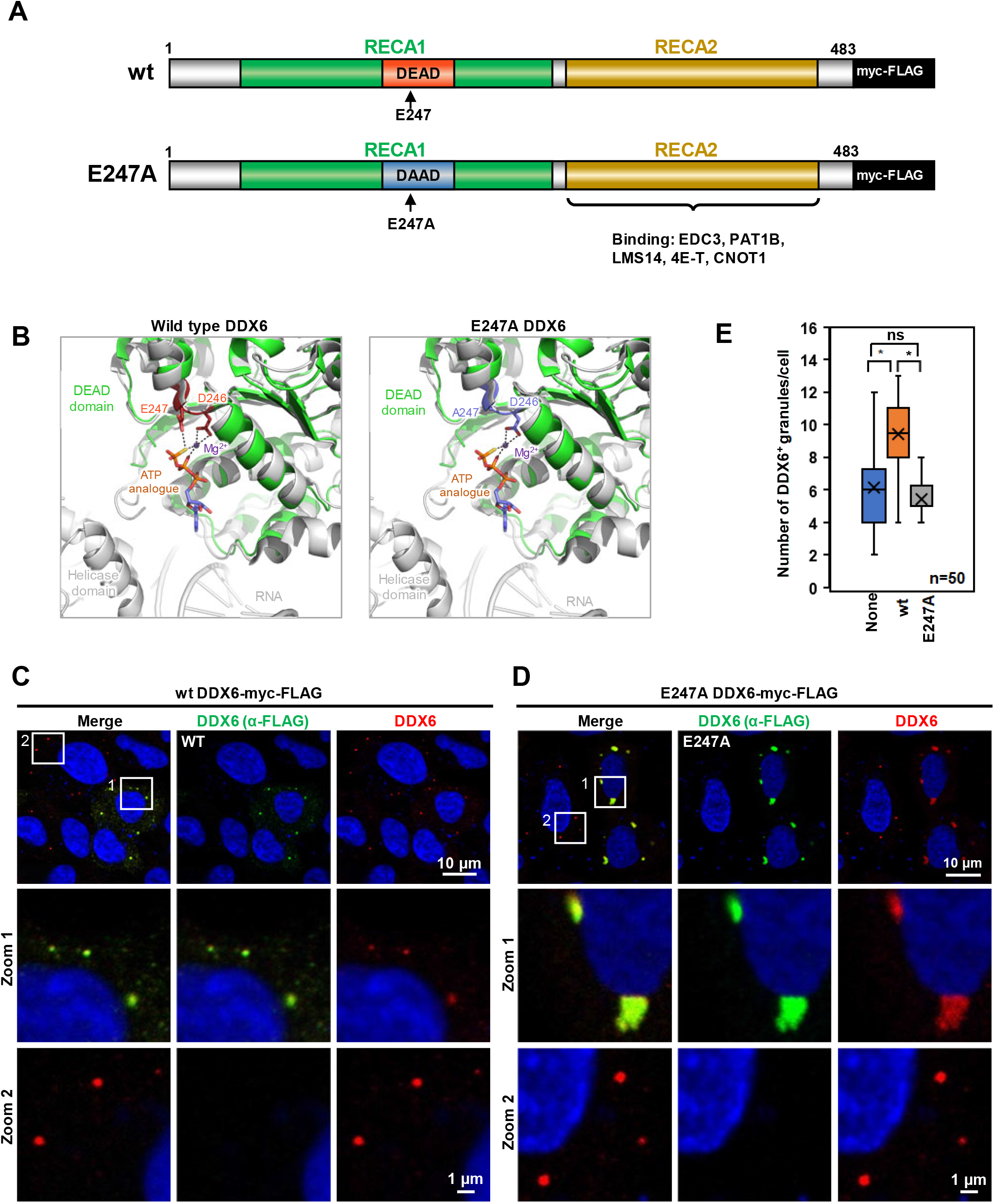
Expression and localization of the ectopic wt DDX6 and helicase-deficient E247A DDX6 mutant. (A) A diagram of ectopically expressed wild type (wt) human DDX6 with two RecA-like domains (RecA1, green, RecA2, yellow). The DEAD-box motif located in the RecA1 is shown in red. The helicase deficient E247A mutant is shown below. Both proteins were fused with c-myc and FLAG tags on the C-terminus. Proteins interacting with the DDX6 C-terminus are also shown. (B) Potential structural basis for the E247A mutation. The DEAD-box motif of DDX6 (PDB ID: 5ANR, green cartoon) is superposed with the DEAD-box of RNA helicase DDX3X at pre-unwound state (PDB ID: 6O5F, gray cartoon) to show the relative locations of DEAD-box and helicase domain. An ATP analogue ATP-γ-S and a Mg^2+^ critical for ATP hydrolysis are modelled based on alignment with the Ski2 RNA-helicase Brr2 (PDB ID: 6QV4). The DEAD-box motif in the wild type and E247A mutant DDX6 are shown in pink and blue, respectively, with the D246 and E/A247 residues in stick representation. Bonds between D246, E247, ATP-γ-S and Mg^2+^ are shown in dash lines. The E247A mutation abolishes critical chemical interactions needed for ATP binding and/or hydrolysis. (C-D) HeLa cells were transfected with wt or helicase-deficient (E247A) DDX6 mutant and stained with anti-FLAG (green) and anti-DDX6 (red) antibodies. (E) HeLa cells were transfected with DDX6 wt or helicase-deficient E247A mutant and were stained with an anti-FLAG antibody or an anti-DDX6 antibody (C and D). Both ectopic and endogenous DDX6^+^ granules in the cells stained with an anti-DDX6 antibody were counted in 50 cells. The average number of the DDX6^+^ granules in 50 cells was plotted in bar graphs. The p-value was calculated by two-tailed Student’s *t*-test (*p<0.001). Ns, no significance.

**Figure 7.**
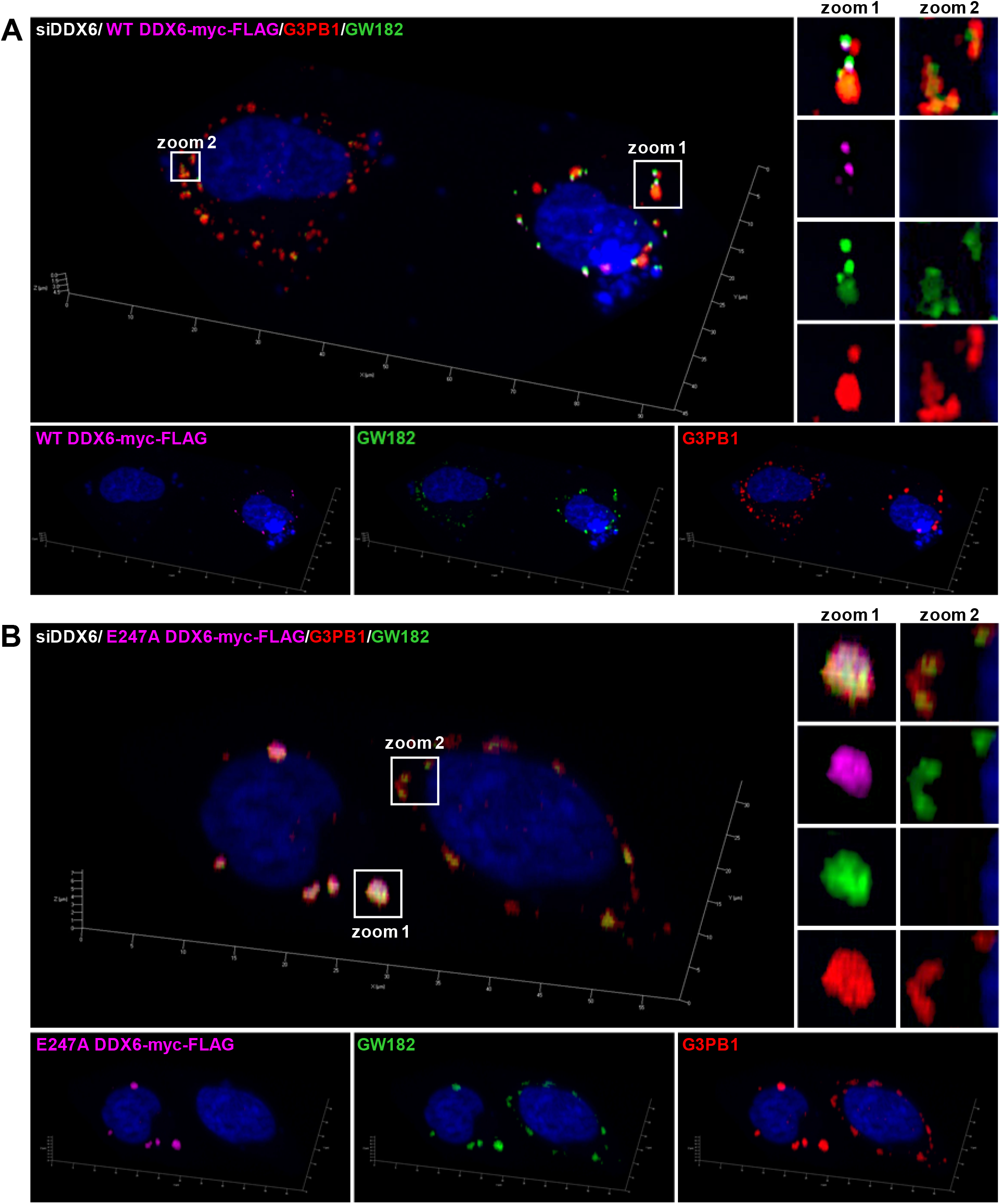
DDX6 wt rescues typical SG formation in cells under ARS stress, but helicase-deficient mutant DDX6 induces SG in cells without ARS stress. HeLa cells were treated twice with DDX6 siRNA in a 24 h interval and then transfected with vectors expressing FLAG-tagged wt (A) or helicase-deficient E247A DDX6 mutant (B). At 24 h after transfection the cells were treated with ARS (0.5 mM for 30 min), fixed, and stained with an anti-FLAG for ectopic DDX6 (magenta), an anti-GW182 for PB (green), and an anti-G3BP1 antibody for SG (red). The cell nuclei are shown in blue. The *z*-stack images in the cells side-by-side with or without ectopic wt DDX6 or E247A DDX6 mutant were collected by confocal microscopy and converted to 3D models using maximum intensity mode.

As shown in **Figure 6C-6D** and **Figure S5**, the wt DDX6-myc-FLAG in HeLa cells form typical PB which can be strained also by both anti-DDX6 and anti-GW182 antibodies under both normal and ARS stress conditions. TIA-1 and PABPC1 showed diffused cytoplasmic staining in the cells without ARS treatment but a distinct punctate pattern of SG in cells with ARS treatment (**Figure S4A**). Despite their overlapping or proximity, PB and SG were readily distinguishable in the cells with ectopic expression of wt DDX6-myc-FLAG (**Figure S4A**). On the other hand, in the cells expressing the helicase-deficient mutant E247A DDX6, the E247A DDX6-positive granules were much bigger in size and also showed strong positivity for both TIA-1 and PABC1 staining surprisedly even in the cell absent of ARS stress (**Figure S4B**). The same was observed after ARS treatment, indicating that they were authentic SG (**Figure S4A**). These data indicated that the DDX6 with deficiency of its helicase activity led to the formation of SG in HeLa cells even under normal physiological condition. Alternatively, the ARS stress provided is presumably toward inhibition of ATPase activity of DDX6 for SG formation in the stressed cells. Colocalization of the E247A DDX6 mutant with SG markers was observed in all investigated cells (N=50) (**Figure S4B**). Consistently, G3BP1 KD could block the SG formation induced by the E247A DDX6, but not PB formation in the cells without or with ARS treatment as seen in the cells with ectopic DDX6 expression (**Figure S5**). Interestingly, similar SG-like G3BP1^+^ granules were also observed in HeLa cells either with overexpression of empty plasmids-derived RNAs or helicase-free GFP-protein, but not in the cells treated only with the transfection reagent used for transfection of the expression vectors (**Figure S6**).

### Normality of regular SG morphology can be restored by ectopic wt DDX6 in the cells lacking endogenous DDX6

Given DDX6 being essential for biogenesis of both PB and SG and to further confirm DDX6 on SG biogenesis, we wish to perform a complementation experiment in the siRNA-mediated DDX6 KD cells to see if loss of the endogenous DDX6 could be rescued by the ectopically expressed exogenous DDX6. As shown in **Figure 6C** and **Figure S7A**, ectopic wt DDX6 formed small, rounded PB in the transfected cells without (**Figure 6C**) or with (**Figure S7A**) non-targeting siRNA siNT treatment, which can be stained by both anti-FLAG, anti-myc and anti-DDX6 (**Figure 6C, Figures S4A, S5**) or anti-GW182 antibodies (**Figure 7A, Figure S5**), whereas the PB in the cells without expression of the ectopic wt DDX6 was stained only by the anti-DDX6 (**Figure 6C**) or anti-GW182 antibody (**Figures S5, S7A**), but not by the anti-FLAG or anti-myc antibody (**Figure 6C, Figures S4, S5**). On average, each transfected cell expressing ectopic wt DDX6 showed slight but significant increase of DDX6^+^ PB number to ∼9.5 granules when compared to the cells expressing only endogenous DDX6 with ∼6 DDX6^+^ PB (**Figure 6E**). In ARS-treated HeLa cells, we found remarkably increased formation of DDX6-FLAG^+^ PB co-stained by GW182 and also the typical shape of SG with TIA-1, PABPC1, and G3BP1 staining mostly overlapping or in proximity to GW182^+^ and DDX6-FLAG^+^ PB (**Figure 7A, Figures S4A, S7A**). These data indicate that the ectopic wt DDX6 acted as and did not interfere with the endogenous DDX6 in the biogenesis of both PB and SG in HeLa cells.

Similar to our previous observations (Sharma et al., 2019), cells with siRNA-mediated KD of DDX6 expression not only blocked PB formation (**Figure S1**), but also displayed the irregular morphology of ARS-induced SG (**Figures 3B, 4B**). However, the DDX6 KD cells with detectable expression of ectopic wt DDX6 exhibited restoration of the formation of GW182^+^ PB under both untreated (**Figure S7A**) and ARS-treated conditions (**Figure 7A, Figure S7A**). More significantly, in the ARS-treated cells with DDX6 KD, the restoration of PB by expression of ectopic wt DDX6 in a cell also reconstituted the morphology of typical G3BP1^+^ SG when compared with a neighboring cell with no wt DDX6 expression (**Figure 7A, Figures S8A, S9**). We also noticed that both ectopic and endogenous DDX6 could contribute to SG biogenesis either separately alone or together (**Figure S9, compare zoom 1 to zoom 2**). These data provide compelling evidence that DDX6 is indeed an essential factor for the biogenesis of both PB and typical SG.

### Helicase-deficient DDX6 mutant functions as a dominant-negative protein on endogenous DDX6 activities

As described above, the helicase-deficient DDX6 mutant induced bigger size of SG co-stained by anti-DDX6 and anti-GW182 in the cells with no ARS stress (**Figure 6D, Figures S5B, S7B**) and sensitive to the KD of G3BP1 in the cells as did with the wt DDX6 (**Figure S5**). Noticeably, it appeared in the E247A DDX6-expressing cells where no endogenous DDX6^+^ or GW182^+^ PB were detectable, which was in contrary to their neighboring cells with no E247A DDX6 expression, but with typical DDX6^+^ or GW182^+^ PB (**Figures 6D, 7B, Figure S7B**). These data indicate that the E247A DDX6 mutant in the cells might function as a dominant-negative protein on endogenous DDX6 activities. Number of DDX6^+^ granules in the E247A DDX6 expressing cells was also fewer (median 5 PB/cell) than that in the cells ectopically expressing the wt DDX6 (median 9.5 PB /cell, **Figure 6E**).

Next, we examined the helicase-deficient E247A DDX6 mutant in the rescue experiment (**Figure 7B, Figures S7B, S8B**). HeLa cells treated with either the non-targeting siNT or DDX6-specific siRNAs were transfected with the E247A DDX6 mutant. Localization of the ectopic E247A DDX6 mutant detected with an anti-FLAG antibody was compared to the localization of endogenous markers for PB (GW182) and SG (G3BP1) in the cells under normal (-ARS) or stress (+ARS) conditions. As shown in **Figure 7B, Figures S7B, S8B**), the E247A DDX6 mutant formed the larger irregular SG similar to the observations in **Figure 6D** and **Figure S4B**, as expected, in the transfected cells even without ARS stress (**Figure S7B**) and had no observed rescuing function. These DDX6 mutant-derived SG were not inducible by ARS stress, while the surrounding neighboring cells with no E247A DDX6 mutant showed the formation of inducible SG in siNT-treated cells or inducible, but irregular SG in the DDX6 KD cells.

## Discussion

A large body of works from many groups have investigated the formation, regulation, and function of PB and SG, but their intimate relation remains elusive. Here, we revisited the dependence of SG biogenesis on PB components in HeLa cells under ARS stress. Our kinetics study observed a topological correlation of SG nucleation to existing PB. We further confirmed the selective role of PB components for SG formation. By siRNA KD of the components required for PB formation in HeLa cells under physiological conditions or ARS stress, we identified DDX6 as a key regulatory factor being required for biogenesis of both PB and consequent SG. Interestingly, except for a minor role of 4E-T and TNRC6A (GW182), the other DDX6 interactors and PB components PAT1B, CNOT1 were found not to contribute to this DDX6 activity.

The role of PB in SG formation and *vice versa* is highly debated and is rather confusing. SG nucleation is regulated by the local concentration of repressed RNAs and SG nucleators. In yeast, the PB were found to promote SG formation (Buchan et al., 2008). The rapid dynamic PB components and partially overlapping proteomes between PB and SG supported this hypothesis (Kedersha et al., 2005; Kroschwald et al., 2015; Youn et al., 2018). The enrichment of specific RBP around PB may perhaps create the sites for the initial step of SG nucleation followed by PB-independent of LLPS and formation of mature SG. Recent studies of *in vitro* SG assembly confirmed the dependence of SG formation on high local concentration of homopolymers in the form of RNA- or synthetic peptides-mediated LLPS by either RNA-RNA or protein-protein interactions (Van Treeck and Parker, 2018). We found that SG are formed topologically in close proximity to or overlapping with PB and contain important PB components, including GW182 and DDX6 (**Figures 1, 2**), consistent with the previous microscopic observations and recent SG proteome analysis (Ayache et al., 2015). More importantly, KD of DDX6 expression leading to deficiency of PB in the cells under physiological and stress conditions and formation of irregular SG in the cells under ARS stress further indicates the importance of DDX6 scaffolding activity in the formation of both PB and SG (**Figure 4B**). An alternative model proposes that translationally arrested RNPs are first targeted to SG, where they are remodeled, and mRNAs stripped of translational components are then transferred to PB for storage and degradation (Mollet et al., 2008; Ohn et al., 2008; Stoecklin and Kedersha, 2013; Youn et al., 2018). However, the inhibition of SG formation did no prevent PB formation (Ohn et al., 2008). We found that KD of G3BP1 expression in HeLa cells in our study did not have any effect on PB formation (**Figure S5**). Others argue that PB and SG are formed independently and PB are “docked” to SG to allow content exchange (Kedersha et al., 2005). Several proteins were identified to promote PB docking on SG such as TTP and CPEB1 (Kedersha et al., 2005; Wilczynska et al., 2005). Nevertheless, a recent finding showing single mRNAs interacting with both SG and PB and moving bi-directionally between SG and PB clearly indicates this intimate relation of PB with SG (Moon et al., 2019).

Our functional studies on PB components required for SG formation identified DDX6 as the major factor necessary for the biogenesis of both PB and SG after ARS treatment (**Figure 3**). DDX6 KD completely abolished the PB formation in cells without ARS treatment (**Figure S1**). The other factors, such as GW182 and 4E-T, were found essential for PB formation under normal physiological conditions too (**Figures S1, S2**), but not much so in ARS-treated cells (**Figures 3, 5**), indicating they are dispensable for PB formation under stress. We also observed that DCP1A was absent in the PB in the 4E-T KD cells under ARS stress (**Figure 5**). The differences in PB formation under normal and stress conditions were observed for other factors (Ayache et al., 2015; Ohn et al., 2008), suggesting the notable changes in PB biogenesis and their compositions during the stress over the physiological condition. In addition, the context- and disease-dependent diversity was observed in SG formation (Markmiller et al., 2018). It will be of great interest to know how the altered protein composition in PB during ARS stress affects the pool of RNAs in PB and what will be a minimal number of essential proteins needed for the formation of PB and subsequent SG.

The dependency of PB formation on DDX6 in normal growth conditions has been well documented from yeast to mammalian cells (Ayache et al., 2015; Buchan et al., 2008). In this report, we now found a significant defect in SG biogenesis in the cells lacking DDX6 (**Figure 4**), where irregular SG induced by ARS stress exhibited less defined shape, lower condensation and overall weaker G3BP1 signal intensity. Strikingly, these atypical SG showed the positivity for numerous PB markers, including DCP1A and GW182 (**Figures 3, 4**). Similar colocalization of PB markers DCP1A or EDC4 with an SG marker eIF3 was observed in retinal pigment epithelial cells and HeLa cells after DDX6 KD (Mollet et al., 2008; Serman et al., 2007). Although the nature of both PB and SG dependency on DDX6 remains to be understood, the observation that the formation of PB and typical SG could be rescued in our study in the cells with DDX6 deficiency by ectopic expression of exogenous wt DDX6 supports the unique functionality of DDX6 in their biogenesis (**Figure 7A, Figure S8**), together with or without endogenous DDX6. However, PB and SG are also enriched with other RNA-helicases, including UPF1, MOV10, eIF4A (DDX2), DDX3 or RNA helicase A (DHX9) (Hubstenberger et al., 2017; Markmiller et al., 2018; Youn et al., 2018). Among these RNA helicases, what and why would be unique for DDX6 to function in the biogenesis of both PB and SG remains unknown. The detailed proteomic and transcriptomic analysis of these two types of RNA granules in the cells with or without DDX6 deficiency under health and ARS stress would shed more light in future to understand its structural and functional correlation to the formation of both PB and SG.

Besides its RNA-binding and helicase activity, DDX6 interacts with a large number of proteins (Ayache et al., 2015). High binding capacity provides DDX6 access to numerous complexes, including translation repressor complex and mRNA decay complexes. These interactions are often mutually exclusive and may have the opposite effect on DDX6 activity (Mugler et al., 2016; Sachdev et al., 2019; Tritschler et al., 2009). DDX6 in cells is highly abundant, with only ∼10% of total DDX6 being localized in PB (Ernoult-Lange et al., 2012). Interestingly, we showed that SG biogenesis is independent of most DDX6-binding partners, except 4E-T, which is required for PB formation under normal conditions as reported (Ayache et al., 2015) but plays a minor role in SG formation under ARS stress (**Figure 5**). Several studies suggested that DDX6 RNA helicase activity was essential for *de novo* PB biogenesis but had no effect on the matured PB (Di Stefano et al., 2019; Minshall et al., 2009; Mugler et al., 2016). We confirmed the importance of DDX6 helicase activity in SG biogenesis by disrupting DDX6 helicase activity by the introduction of an E-to-A mutation into the DEAD-box. Structural modeling with other DEAD box helicase, such as the DDX3 (Song and Ji, 2019) and Ski2 RNA-helicase Brr2 (Absmeier et al., 2020), indicated that the DDX6 E247 may be critical in mediating ATP-binding and hydrolysis that is needed for DDX6 function (**Figure 6B**). We found this functional deficient mutant E247A DDX6, but not the wt DDX6 in transfected cells, induces the formation of SG positive for DDX6 and GW182 staining under a stress-free condition (**Figure 6D, Figures S4B, S7B**). Since ATP binding and hydrolysis is important for DDX6 helicase activity to function, the mutation in E247A DDX6 will reduces ATP binding and ATP hydrolysis (**Figure 6B**), thus, mimicking the effect of ARS which blocks the production of ATP (Finnegan and Chen, 2012; Jain et al., 2016) being required for DDX6 activity. Although the exact machinery how the E247A DDX6 mutant induces SG-like granules or its formation resembles the type II SG formation (Bordeleau et al., 2006; Dang et al., 2006) in the transfected cells remains to be determined, interestingly, we found under a stress-free condition that HeLa cells expressing empty vector-derived RNAs or even helicase-free GFP also exhibited these SG-like granules.

Together, these data clearly indicate that DDX6 helicase activity is needed to prevent SG formation while is essential for PB biogenesis in cells under physiological condition. Stress conditions provide the cellular machinery to block the RNA helicase activity of DDX6 and thereby promote SG formation. In this regard, DDX6 functionally resembles eIF4A, another SG-associated DEAD-box RNA-helicase (Tauber et al., 2020), in prevention of SG formation in the cells. Thus, our study provides a compelling evidence demonstrating the opposite roles of DDX6 RNA helicase in the biogenesis of both PB and SG.

## Material and methods

### Cells

HeLa cells (ATCC, Manassas, VA) were maintained in Dulbecco’s Modified Eagle Medium (DMEM, Thermo Fisher Scientific, Waltham, MA) supplemented with 10% fetal bovine serum (FBS, Hyclone, Cytiva, Marlborough, MA) and 1 × Penicillin-Streptomycin-Glutamine (Thermo Fisher Scientific). The cells were kept at 37°C in a humidified atmosphere containing 5% CO_2_.

### Stress induction

HeLa cells were incubated in the complete culture medium supplemented with 0.5 mM sodium arsenite (ARS, Sigma Aldrich, St. Louis, MO) for 0-30 min to induce the stress granule formation. Cells without ARS treatment were used as a negative control.

### siRNA knockdown

HeLa cells at 0.25 × 10^6^ were plated into 6-well plates 24 h before the first siRNA transfection with 40 nM of the ON-TARGETplus human siRNA SMARTpool (Horizon Discovery Biosciences) using LipoJet (SignaGen Laboratories, Frederick, MD) as recommended by the manufacturer. Twenty-four hours after the siRNA transfection, the cells were split into two wells with glass coverslips and four hours later, transfected again with the same siRNA at 40 nM, followed by additional 24 h incubation before the stress induction. ON-TARGETplus human siRNA SMARTpools (Horizon Discovery Biosciences) targeting DCP1A (L-021242-00-0005), DDX6 (L-006371-00-0005), EIF4ENIF1 (4E-T, L-013237-01-0005), PAT1B (L-015591-01-0005), G3BP1 (L-012099-00-0005), CNOT1 (L-015369-01-0005), TIA-1 (L-013042-02-0005), TNRC6A (L-014107-00-0005) were used in this study. An ON-TARGET plus Non-targeting siRNA #1 (siNT, D-001810-01-05, Horizon) served as a negative control.

### Complementation assay

The vector expressing myc-DDK-tagged-human wt DDX6 was purchased from OriGene (RC209431, Rockville, MD). The helicase-deficient DDX6 E247A mutant was constructed by overlapping PCR using primers oVM506 5’-ACTTATCTGCCGCATCCAATACTATCATCTGGACATGAT-3’ and oVM507 5’-TTGGATGCGGCAGATAAGTTGCTGTCACAGGATTTTGTG-3’. HeLa cells were treated with non-targeting (siNT) or DDX6-targeting (siDDX6) siRNA twice at a 24 h interval. At two hours after the second siRNA transfection, the cells were transfected with the vector expressing either a wt or E247A mutant DDX6. After 24 h of vector transfection, the HeLa cells were then subjected to 0.5 mM ARS treatment for 30 min before immunofluorescent staining.

### Immunofluorescent (IF) staining

HeLa cells growing on a cover glass (-/+ ARS) were fixed with 4% paraformaldehyde (Electron Microscopy Sciences, Hatfield, PA) in 1 × PBS (phosphate buffered saline, pH 7.4) for 20 min at RT, followed by 5 min quenching with 100 mM glycine and permeabilization with 0.5% Triton X-100 (Promega, Madison, WI) in 1 × PBS for 15 min. The cells were then washed three times with 1 × PBS and blocked with 3% Blot-Qualified bovine serum albumin (BSA, Promega) in 1 × PBS containing 0.05% Tween-20 (TPBS) in a humidified chamber for 1 h at 37°C. The primary (1:100) and AlexaFluor-labeled secondary antibodies (1:500, Thermo Fisher Scientific) were diluted in blocking solution and incubated with the slides in a humidified chamber at 37°C for 1-2 h. After each incubation, the slides were washed three times with TPBS. Hoechst 33342 (Thermo Fisher Scientific) was added to the last wash (1 µg/ml) to stain cell nuclei for 5 min. The fully stained slides were mounted using ProLong Gold Antifade Mountant (Thermo Fisher Scientific) and let cure overnight at RT.

### Image collection and processing

The IF-labeled HeLa cells were examined with Zeiss 710 confocal microscope using 63 × oil objective. The pictures were collected and processed using ZEN imaging software (Zeiss). The 3D images were generated from respective *z*-stack images using the maximum intensity function. The ImageJ (https://imagej.nih.gov) was used to measure the signal intensity for individual channels at the cross-sections.

### Antibodies

The following primary antibodies were used in the study: human autoimmune serum containing anti-GW182 antibody were obtained as gifts from Dr. Marvin Fritz, University of Calgary; others were anti-DCP1A (ab183709, Abcam, Cambridge, MA), anti-TIA-1 (OTI1D7 clone)(MA5-26474, Thermo Fisher Scientific), anti-G3PB1 (611126, BD Biosciences), anti-PABPC1 (ab21060, Abcam), anti-EDC3 (16486-1-AP, Proteintech, Rosemont, IL), anti-DDX6 (NB200-191, Novus Biologicals, Centennial, CO), anti-CNOT1 (14276-1-AP, Proteintech), anti-PAT1B (A303-482A, Bethyl Laboratories, Montgomery, TX), anti-β-tubulin (T5201, Millipore Sigma), rabbit anti-FLAG (F7425, Millipore Sigma) and mouse anti-c-Myc−FITC antibody (F2047, Millipore Sigma).

### Western blot

The cells transfected twice with non-targeting (siNT) or gene-specific siRNAs at a 24 h interval were harvested by direct lysis in 2 × SDS protein samples buffer containing 5% (vol/vol) 2-mercaptoethanol. The obtained protein samples were separated on 4-12% Bis-TRIS NuPAGE gel in 1 × MES buffer (Thermo Fisher Scientific) and transfer to nitrocellulose membrane before blocking with 5% non-fat milk dissolved in 1 × TRIS-buffered saline (TBS) containing 0.05% Tween-20 (TTBS). The membranes were stained with primary antibodies diluted 1:1,000 in TTBS followed by incubation with horse peroxidase-labeled secondary antibody (Millipore Sigma) diluted 1:10,000 in 2% non-fat milk in TTBS. The signal was developed by ProSignal Pico ECL Reagent (GeneSee Scientific, San Diego, CA) and processed by ChemiDoc Touch molecular imager (BioRad, Hercules, CA).

### Structural modeling

The crystal structure of DDX6 (PDB ID: 5ANR) (Ozgur et al., 2015) was used to analyze the structural basis of E247A mutation. Briefly, the domain 1 (DEAD domain) of DDX6 was superposed with the corresponding domain of RNA helicase DDX3X at pre-unwound state (PDB ID: 6O5F) (Song and Ji, 2019) to show the relative locations of the DEAD domain and helicase domain with RNA bound. Since the DEAD sequence is a Walker B motif that has been shown to be critical in binding and hydrolysis of ATP, a non-hydrolysable ATP analogue, ATP-γ-S, and a Mg^2+^ critical for ATP hydrolysis were modelled into the DDX6 ATP-binding site based on alignment with the crystal structure of Ski2 RNA-helicase Brr2 (PDB ID: 6QV4) (Absmeier et al., 2020). Structural alignment, modelling and graphical rendering were carried out with PyMol Molecular Graphics System, version 2.4.1 (www.pymol.org).

## Supplementary Information

**Figure S1.**
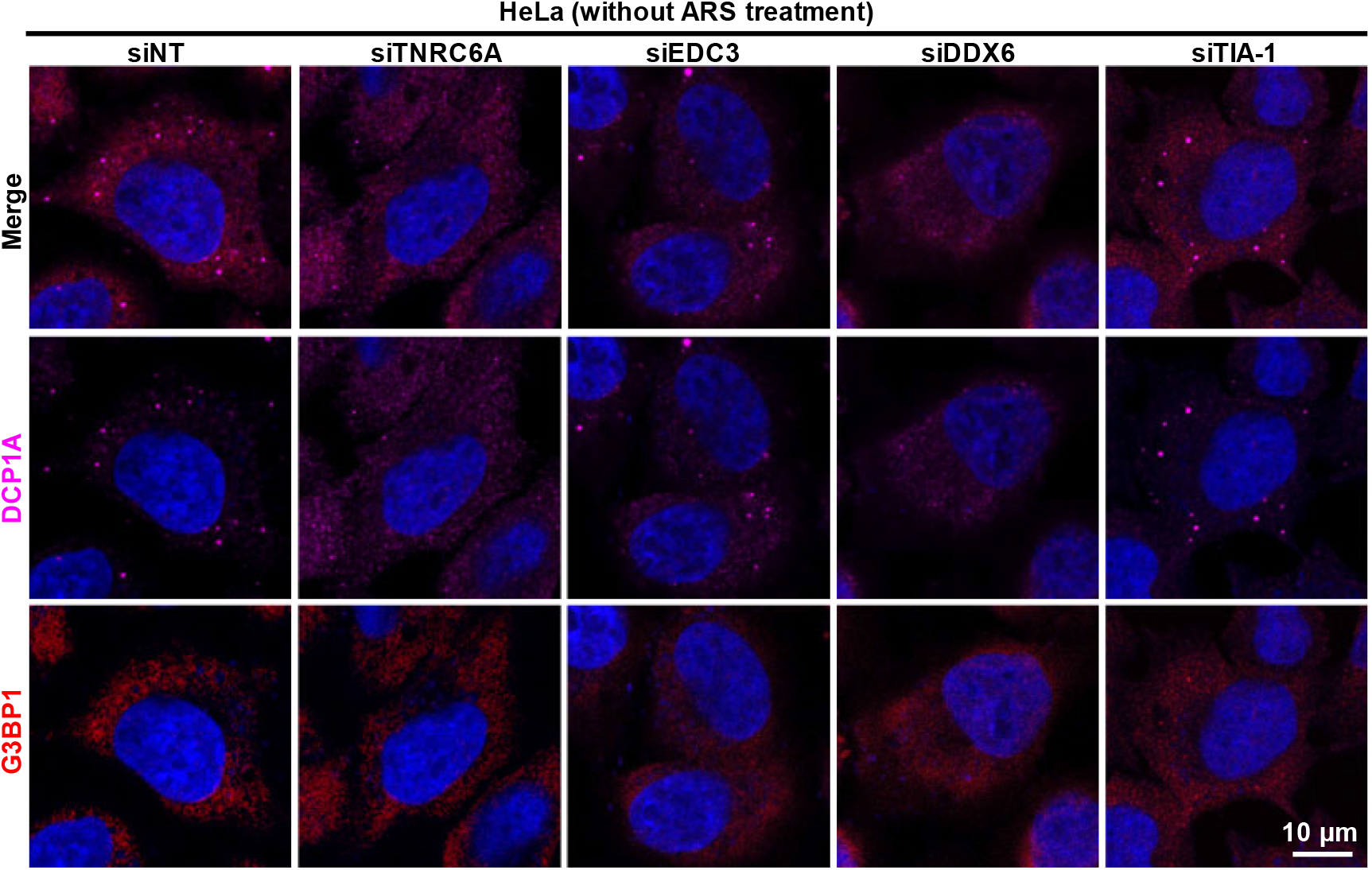
Effect of TNRC6A (GW182), EDC3, DDX6, or TIA-1 knockdown on PB formation in HeLa cells. HeLa cells were treated twice with control non-targeting (siNT) or gene-specific siRNAs and stained with an anti-DCP1A for PB (magenta) and an anti-G3BP1 antibody for SG (red), respectively. Cell nuclei (blue) were counterstained with Hoechst 33342.

**Figure S2.**
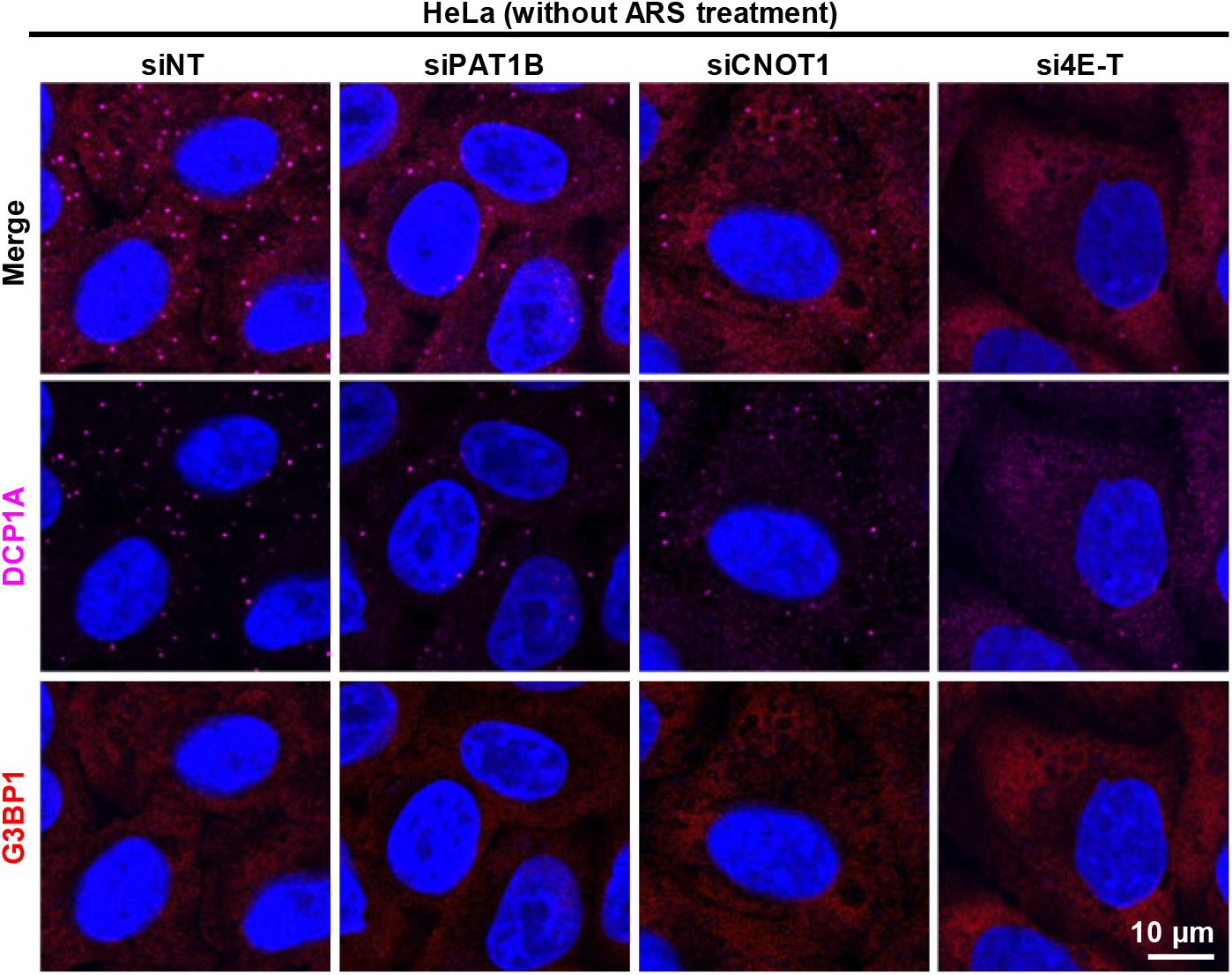
Effect of PAT1B, CNOT1 or 4E-T knockdown on PB formation in HeLa cells. HeLa cells were treated twice with control nontargeting (siNT) or gene-specific siRNAs and stained with an anti-DCP1A for PB (magenta) and an anti-G3BP1 antibody for SG (red), respectively. Cell nuclei (blue) were counterstained with Hoechst 33342.

**Figure S3.**
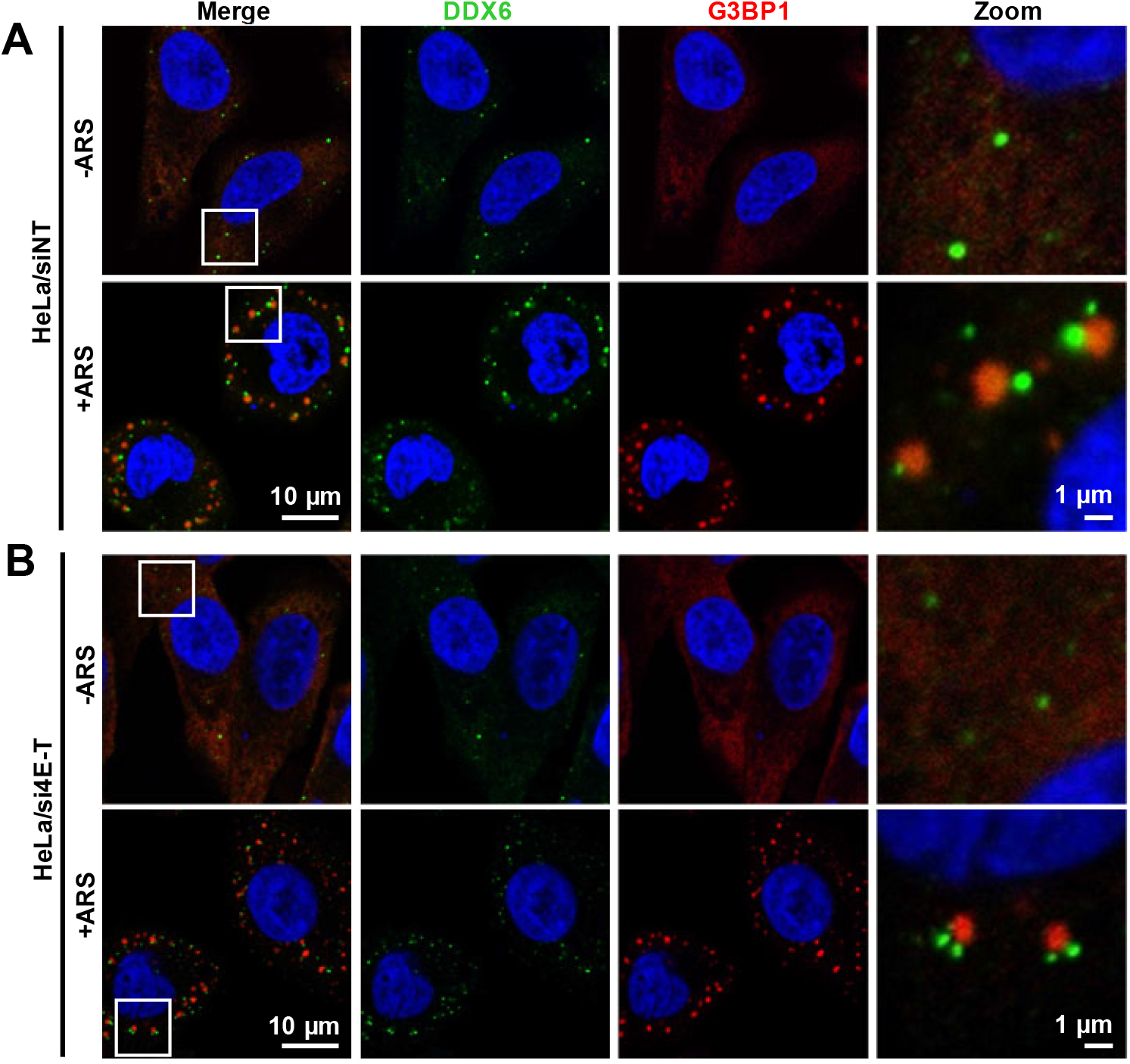
4E-T is dispensable for DDX6 recruitment to PB during the stress. HeLa cells were treated twice with a non-targeting (siNT) (A) or 4E-T targeting (B) siRNA. The cells were then treated without or with ARS (0.5 mM 30 min), fixed and stained for PB marker DDX6 (green) and SG marker G3BP1 (red). Cell nuclei (blue) were counterstained with Hoechst 33342.

**Figure S4.**
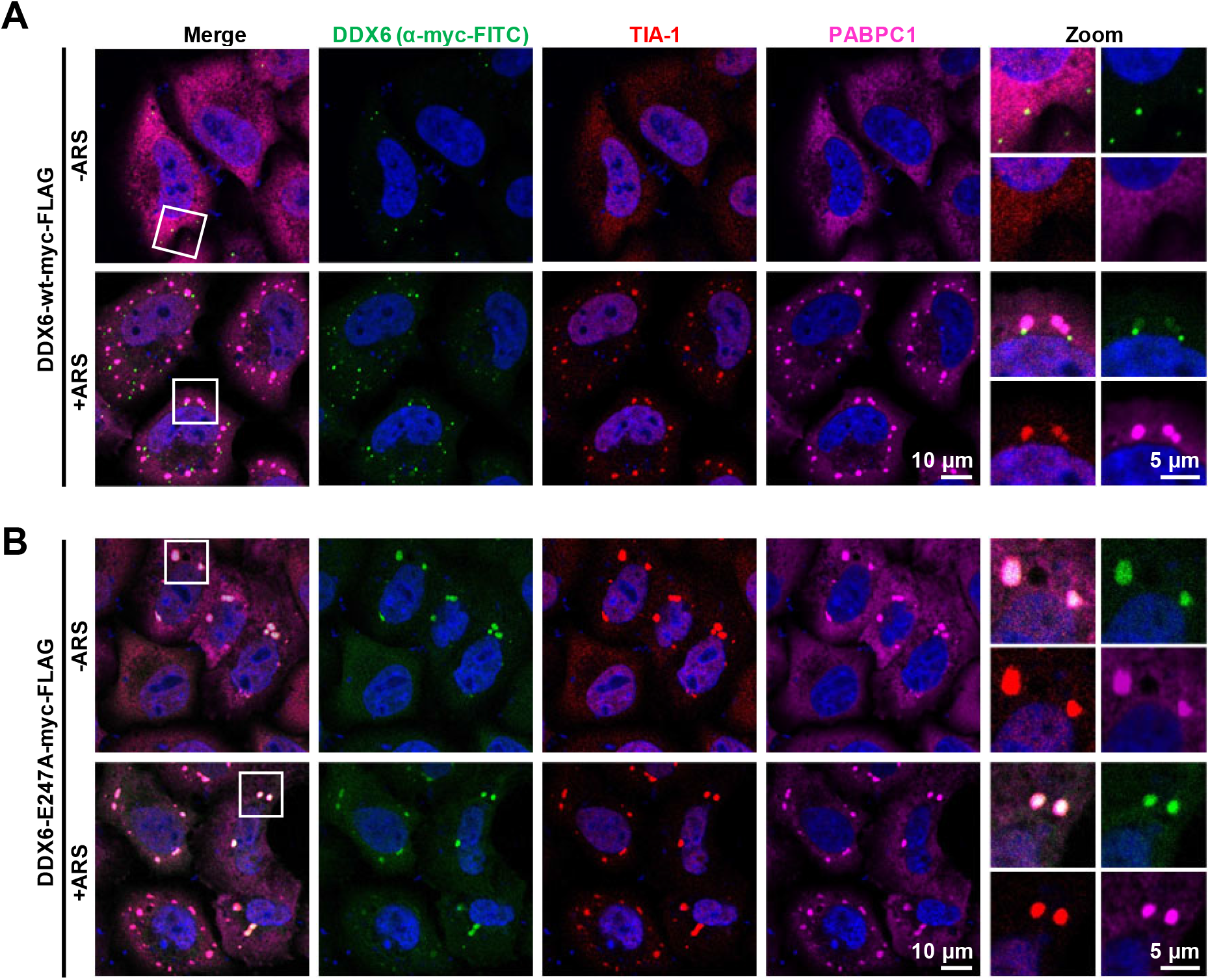
The granules induced by helicase-deficient mutant resemble SG. HeLa cells were transfected with the vectors expressing the myc-FLAG-tagged wt and helicase-deficient (E247A) DDX6 mutant. At 24 h after the transfection ARS (0.5 mM) was added to one half of the samples and incubated for 30 min to induce the SG. Both samples without (-ARS) or with (+ARS) ARS treatment were stained with an anti-myc-FITC for ectopic DDX6 (green), and an anti-TIA-1 (red) and an anti-PABPC1 antibody (magenta) for SG. Cell nuclei (blue) were counterstained with Hoechst 33342.

**Figure S5.**
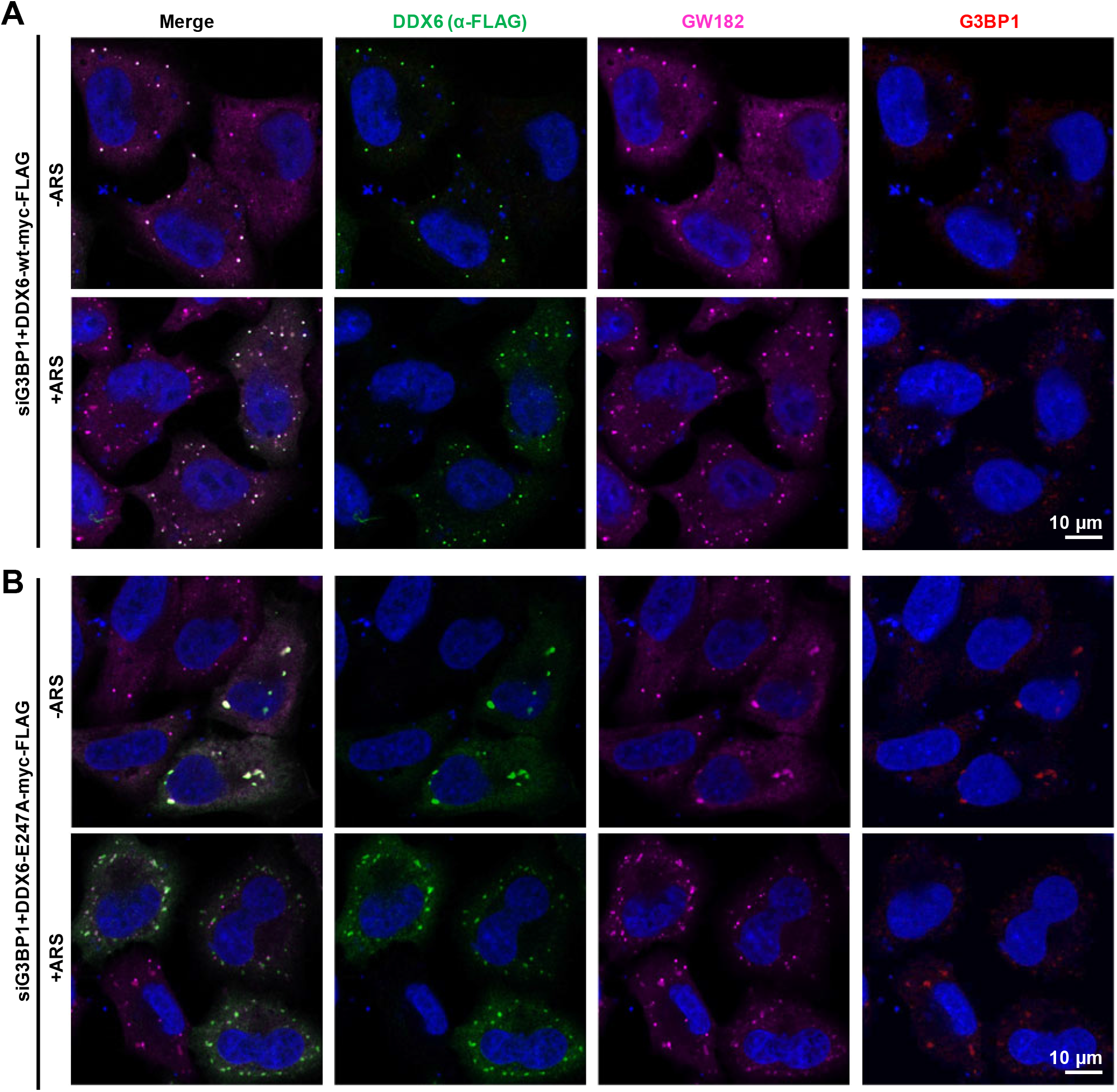
Effect of G3BP1 knockdown on SG induced by DDX6 helicase-deficient mutant. HeLa cells were treated twice with G3BP1 siRNA and then transfected with vectors expressing myc-FLAG-tagged wt or helicase-deficient DDX6 mutant (E247A). At 24 h after the transfection one half of the samples was treated with ARS (0.5 mM) to induce SG. Both samples without (-ARS) or with (+ARS) ARS treatment were stained with an anti-FLAG to detect the ectopic DDX6 protein (green), an anti-GW182 for PB (magenta), and an anti-G3BP1 antibody for SG (red). Cell nuclei (blue) were counterstained with Hoechst 33342.

**Figure S6.**
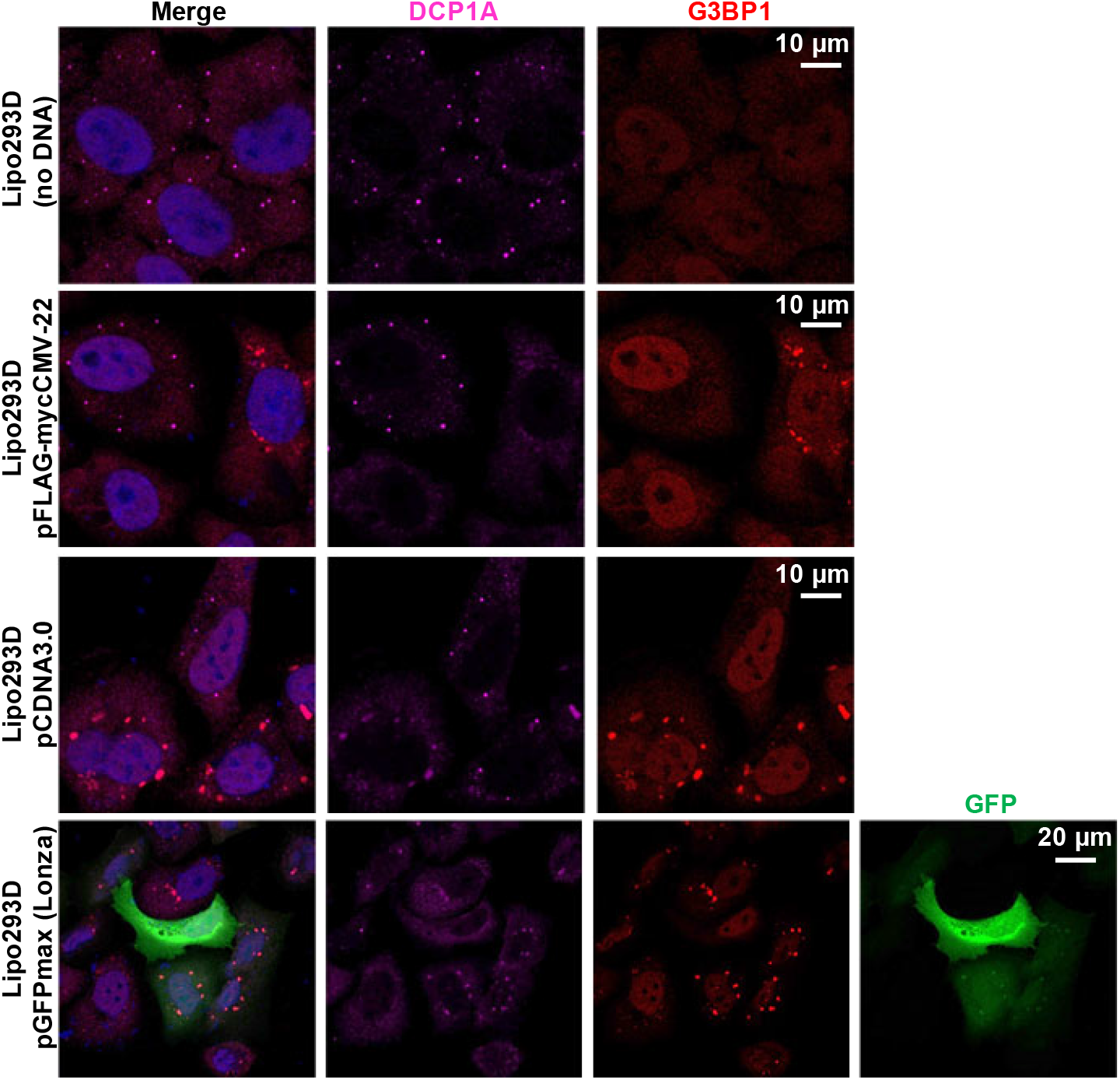
The SG-like granules induced by plasmid-derived RNA in transfected cells with no ARS treatment. HeLa cells were transfected with the empty pcDNA3 (Invitrogen), pFLAG-myc-CMV-22 (Sigma-Aldrich) or GFP expressing pGFPmax (Lonza) vectors using LipoD293 In Vitro DNA Transfection Reagent (SignaGen Laboratories). The cell treated with the transfection reagent alone without any DNA were used as a negative control. At 24 h after the transfection the cells were fixed and stained for PB marker DCP1A (magenta) and SG marker G3BP1 (red). The GFP expression (green) was determined by direct fluorescence. Cell nuclei (blue) were counterstained with Hoechst 33342.

**Figure S7.**
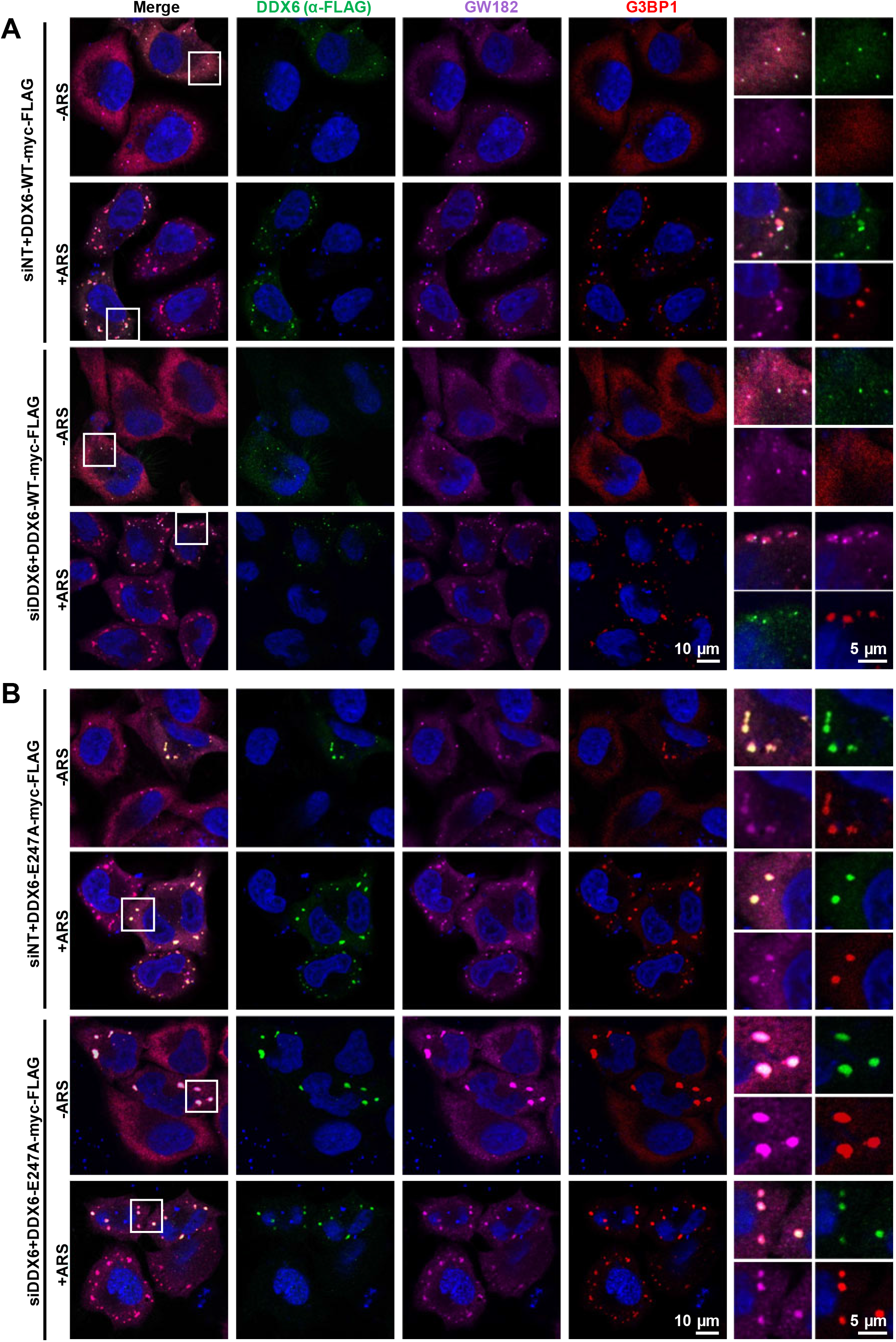
DDX6 helicase-deficient mutant, but not DDX6 wt, induces SG formation in ARS stress-free HeLa cells with or without KD of endogenous DDX6. HeLa cells were treated twice with non-targeting (siNT) or DDX6-targeting siRNA (siDDX6) in a 24 h interval and the transfected with vectors expressing FLAG-tagged wt (A) or helicase-deficient (E247A) DDX6 mutant tagged with myc-FLAG (B). At 24 h after vector transfection the cells were treated without or with ARS (0.5 mM for 30 min), fixed and stained with an anti-FLAG for ectopic DDX6 (green), an anti-GW182 for PB (magenta) and an anti-G3BP1 antibody for SG (red). Cell nuclei (blue) were counterstained with Hoechst 33342.

**Figure S8.**
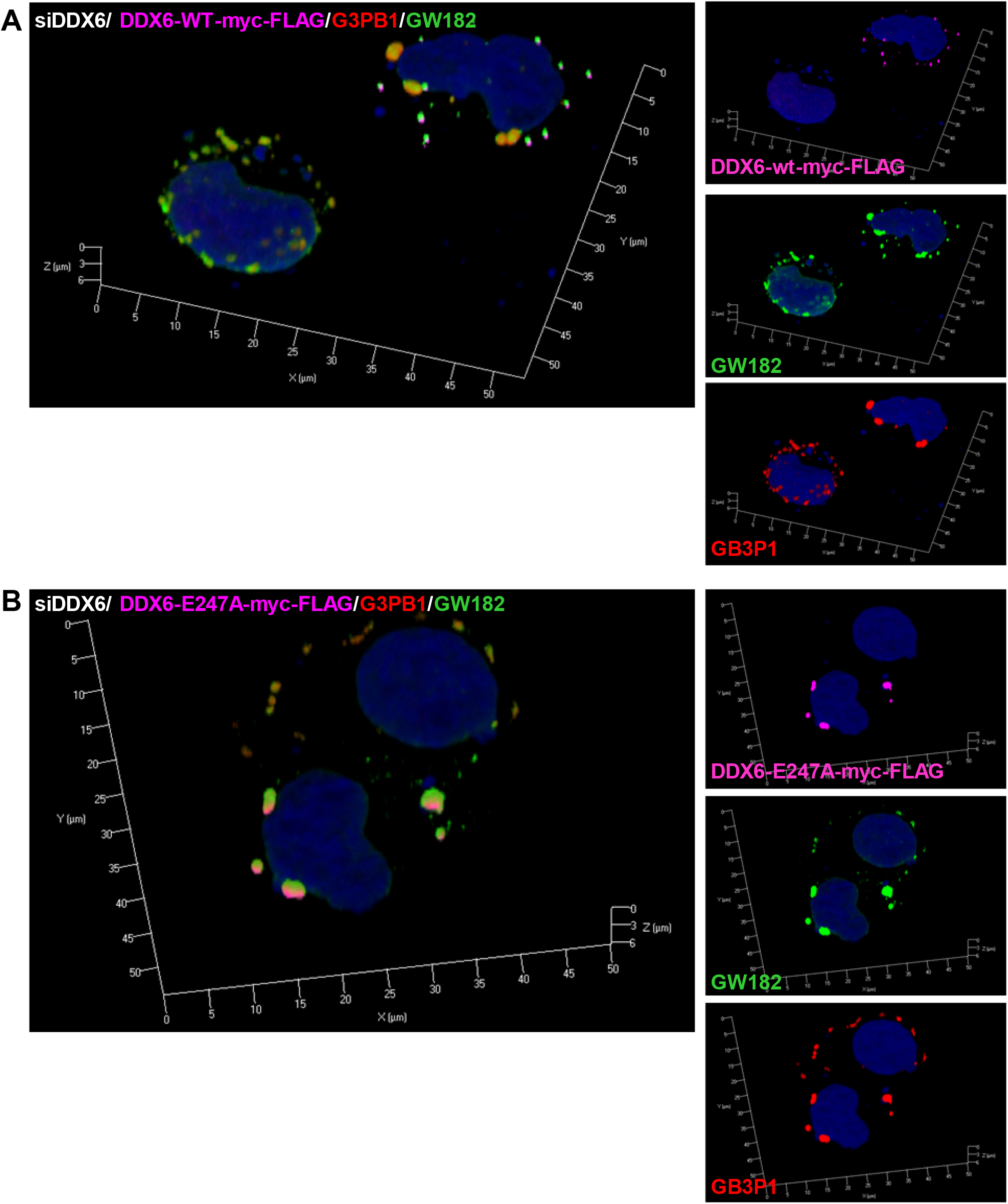
Shape comparison of the enlarged SG in ARS-stressed cells without endogenous DDX6 to the SG in the cells expressing ectopic wt DDX6 or E247A DDX6 mutant. HeLa cells were treated twice with a DDX6-specific siRNA in a 24 h interval and then transfected with vectors expressing FLAG-tagged wt (A) or helicase-deficient (E247A) DDX6 mutant (B). At 24 h after vector transfection the cells were treated with ARS (0.5 mM for 30 min), fixed and stained with an anti-FLAG for ectopic DDX6, an anti-GW182 for PB, and an anti-G3BP1 antibody for SG. The maximum intensity 3D images were generated from respective z-stack images using Zen software.

**Figure S9.**
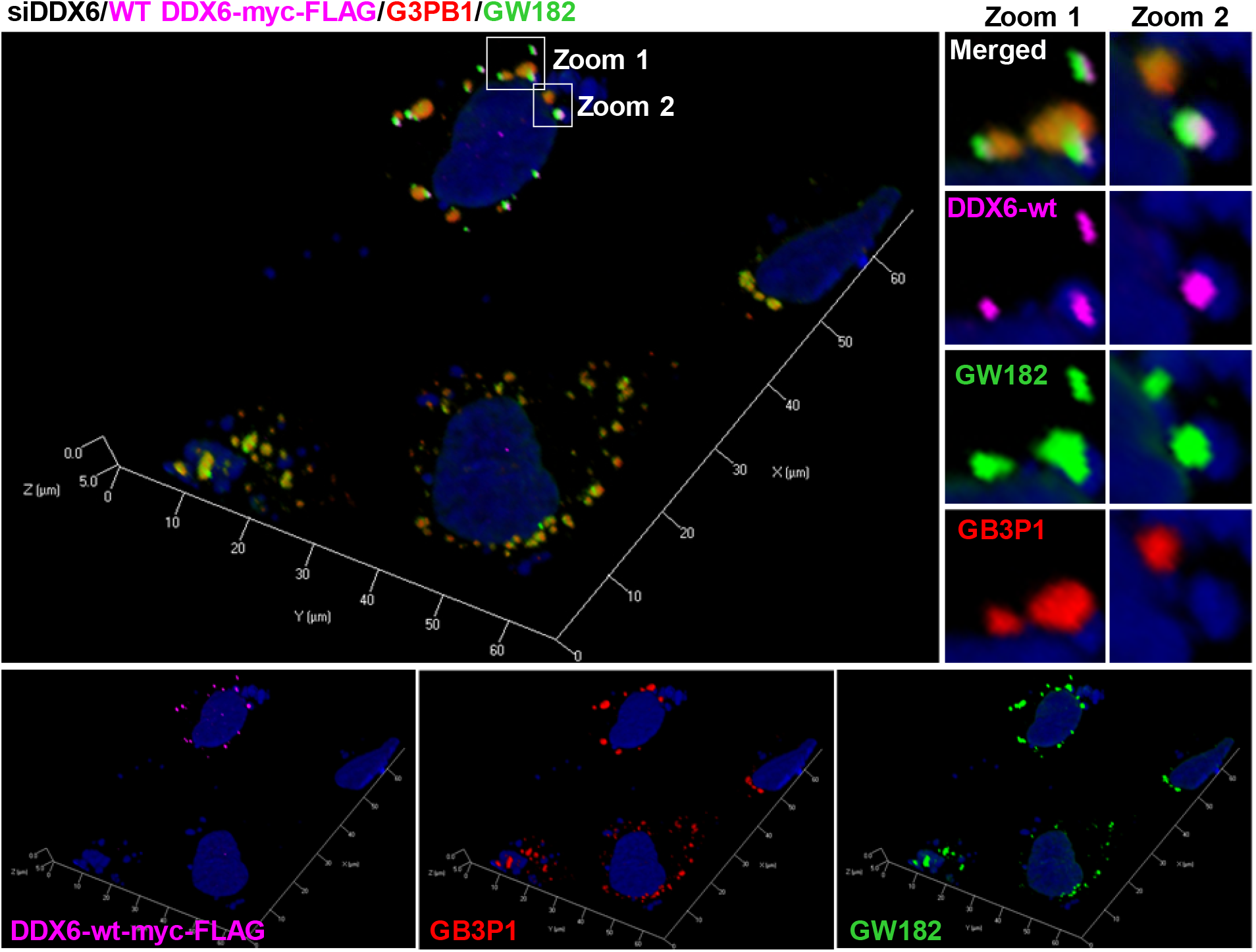
Contribution of both exogenous and endogenous DDX6 to SG formation. HeLa cells were treated twice with a DDX6-specific siRNA (siDDX6) in a 24 h interval and then transfected with vectors expressing myc-FLAG-tagged wt DDX6. At 24 h after vector transfection the cells were treated with ARS (0.5 mM for 30 min), fixed and stained with an anti-FLAG for ectopic DDX6 (magenta), an anti-GW182 for PB (green), and an anti-G3BP1 antibody for SG (red). The nuclei (blue) were counterstained with Hoechst 33342 staining. The maximum intensity 3D images were generated from respective *z*-stack images using Zen software (Zeiss).

## Acknowledgment

We want to thank Dr. Marvin Fritzler of the University of Calgary for kindly providing the autoimmune serum for GW182 staining. We would also gratefully acknowledge the Optical Microscopy and Analysis Laboratory of the Frederick National Laboratory for Cancer Research for their scientific support.

## Authors’ contributions

VM and ZMZ conceptualized, and VM performed the experiments. TZ performed DDX6 structural modeling analysis. VM and ZMZ wrote the manuscript. All authors finalized and approved the manuscript submission.

## Ethics Statement

No human subjects or animals were used in this study.

## Consent for publication

Consent for publication was obtained from all participants.

## Authors’ details

VM and ZMZ: Tumor Virus RNA Biology Section, HIV Dynamics and Replication Program, National Cancer Institute (NCI), National Institutes of Health, 1050 Boyles Street, Bldg. 560, Rm. 11-2, Frederick, MD, 21702, USA

TZ: Structural Biology Section, Vaccine Research Center, National Institute of Allergy and Infectious Diseases (NIAID), National Institutes of Health, 40 Convent Drive, Bldg. 40, Rm. 4609B, MSC3027, Bethesda, MD, 20892, USA

